# Internal states as a source of subject-dependent movement variability and their representation by large-scale networks

**DOI:** 10.1101/2022.08.16.504130

**Authors:** Macauley Smith Breault, Pierre Sacré, Zachary B. Fitzgerald, John T. Gale, Kathleen E. Cullen, Jorge A. González-Martínez, Sridevi V. Sarma

## Abstract

A human’s ability to adapt and learn relies on reflecting on past performance. Such reflections form latent factors called internal states that induce variability of movement and behavior to improve performance. Internal states are critical for survival, yet their temporal dynamics and neural substrates are less understood. Here, we link internal states with motor performance and neural activity using state-space models and local field potentials captured from depth electrodes in over 100 brain regions. Ten human subjects performed a goal-directed center-out reaching task with perturbations applied to random trials, causing subjects to fail goals and reflect on their performance. Using computational methods, we identified two internal states, indicating that subjects kept track of past errors and perturbations, that predicted variability in reaction times and speed errors. These states granted access to latent information indicative of how subjects strategize learning from trial history, impacting their overall performance. We further found that large-scale brain networks differentially encoded these internal states. The dorsal attention network encoded past errors in frequencies above 100 Hz, suggesting a role in modulating attention based on tracking recent performance in working memory. The default network encoded past perturbations in frequencies below 15 Hz, suggesting a role in achieving robust performance in an uncertain environment. Moreover, these networks more strongly encoded internal states and were more functionally connected in higher performing subjects, whose learning strategy was to respond by countering with behavior that opposed accumulating error. Taken together, our findings suggest large-scale brain networks as a neural basis of strategy. These networks regulate movement variability, through internal states, to improve motor performance.

**Key points:**

- Movement variability is a purposeful process conjured up by the brain to enable adaptation and learning, both of which are necessary for survival.
- The culmination of recent experiences—collectively referred to as internal states—have been implicated in variability during motor and behavioral tasks.
- To investigate the utility and neural basis of internal states during motor control, we estimated two latent internal states using state-space representation that modeled motor behavior during a goal-directed center-out reaching task in humans with simultaneous whole-brain recordings from intracranial depth electrodes.
- We show that including these states—based on error and environment uncertainty—improves the predictability of subject-specific variable motor behavior and reveals latent information related to task performance and learning strategies where top performers counter error scaled by trial history while bottom performers maintain error tendencies.
- We further show that these states are encoded by the large-scale brain networks known as the dorsal attention network and default network in frequencies above 100 Hz and below 15 Hz but found neural differences between subjects where network activity closely modulates with states and exhibits stronger functional connectivity for top performers.
- Our findings suggest the involvement in large-scale brain networks as a neural basis of motor strategy that orchestrates movement variability to improve motor performance.

## Introduction

Professional athletes represent the pinnacle of motor control and precision, but they too fall victim to slight variations in their movement. Movement variability is traditionally viewed as a byproduct of noise accumulated by the motor system^[1]^. However, there is emerging evidence that this variability is purposefully orchestrated by the brain to facilitate learning and adaptation^[2, 3]^. For example, more movement variability through exploration leads to faster learning and better performance^[4, 5]^. The decision to explore—as opposed to exploit—an environment to gather information to inform future behavior through learning lends itself naturally to movement variability. Not only does this information depend on the present, but also on internalized factors that account for the accumulation of past experience. These factors are called *internal states*. For example, movement variability is influenced by motivation^[6–8]^, confidence^[9, 10]^, and emotion^[11–13]^. Future behavior is the culmination of current information and internal states.

With their impact on behavior so apparent, it is surprising how ambiguous internal states are in motor control compared to other fields such as decision-making. To date, research into decision-making has used statistical models to explore relationships between behavioral variability and internal states^[14–17]^ with the aim of finding evidence of the brain encoding states related to uncertainty^[18]^, bias^[19]^, trial history^[20, 21]^, and impulsiveness^[22]^. Like decision-making, the goal of motor control is to produce actions that optimize outcomes in the presence of uncertainty^[23]^. Whether those actions be decisions or movements, variability and internal states are inherent to both. Therefore, we speculated that internal states are encoded in regions that are not specific to motor control (*i.e.*, nonmotor regions). Indeed, decision-making tasks that require movements find that regions involved with sensorimotor integration encode their internal states as opposed to motor regions^[19, 20]^. In this context, an emerging consensus is that movement variability originates from both the planning^[24]^ and execution phases^[25]^ of movement. However, the gap in our understanding of motor control becomes apparent when one asks how internal states evolve, where they are encoded in the brain, and how they affect performance.

Two challenges need to be overcome to address these questions. The first challenge is determining the internal states based on behavior. To date, direct measures of the brain’s internal states have remained elusive^[26]^. Researchers have tried to capture measures of such states using methods including self-reporting^[27]^, galvanic skin conductance^[28]^, heart rate variability^[29, 30]^, and pupil size^[31]^. However, these measures are context-dependent^[32]^, vary between individuals^[30]^, and can function with timescales on the order of minutes to compute^[28, 30]^, whereas internal states can change within seconds^[33]^. By comparison, decision-making studies rely on observable behavior such as reaction times, decisions, and outcomes to derive their internal states. Therefore, motor control studies would also be ideal for deriving internal states due to their abundant movement-related data. Even so, since these methods are all byproducts of the brain, the question arises why not measure internal states directly from the brain?

This leads to our second challenge, which is identifying where internal states are encoded in the brain. As previously mentioned, research on decision-making supports the view that internal states are encoded by diverse brain regions involving multiple systems (*e.g.*, sensory and memory)^[18–22]^. Whole-brain imaging with high temporal resolution would be ideal to capture diverse brain structures and rapidly evolving internal states. Most work in humans has used non-invasive neural imaging, such as functional Magnetic Resonance Imaging (fMRI) studies. However, the limitations of the temporal resolution of fMRI^[34]^, compounded by the limited space that subjects must perform a natural movement, make it difficult to link neural correlates of internal states to behavior. What is needed is millisecond resolution with whole-brain coverage.

To address these challenges, we combined high-quality measurements of natural reaching movements with high-spatial and temporal resolution neural StereoElectroEncephaloGraphy (SEEG) recordings. Specifically, ten human subjects implanted with intracranial depth electrodes performed a simple motor task that elicited movement variability during planning and execution. We estimated two internal states using state-space models using measurable behavioral data: the “error state” accumulates based on past errors to convey overall performance and the “perturbed state” accumulates based on past perturbations to convey environmental uncertainty. Adding these states improved our ability to estimate trial-by-trial reaction times and speed errors over using stimuli alone. Our approach also granted us access to latent terms that predicted subject performance and provided insight into subject strategy. Remarkably, we found neural evidence of the brain encoding each of the internal states in relatively distinct large-scale brain networks. Specifically, the Dorsal Attention Network (DAN) and Default Network (DN) were linked to the error and perturbed state, respectively. We also linked the encoding strength and functional connectivity of these networks back to subject performance and strategy.

## Results

To investigate the coupling between motor variability and internal states, we devised a goal-directed reaching task that elicited movement variability both within and between human subjects. We first characterized this variability for our population of subjects during both planning and execution based on trial conditions. Then, to account for this variability, we built a simple behavioral model that incorporated dynamic internal states as accumulating trial history that evolve over time. Using computational methods, we used this model to predict differences in strategy across subjects by splitting subjects into top and bottom performers and compared how these groups use internal states to inform future behavior. Finally, using recordings from intracranial depth electrodes implanted in the same subjects, we investigated the relationship between neural activity in large-scale brain networks and the encoding of these internal states.

### Motor task produced movement variability within and between subjects

Subjects were instructed to perform a motor task in which they made reaching movements toward a target at an instructed speed with the possibility of physical perturbations for a virtual monetary reward using a robotic manipulandum. As visually shown in Figure 1a, this task elicited movement variability. We quantified this variability by calculating the Reaction Times (RT) and Speed Errors (SE) for planning and execution phases, respectively. Figure 1b shows the RT (top) and SE (bottom) results for each subject overlaid across trials, with an example subject (subject 6) traced in black. Both measurements were inconstant across trials for subject 6, confirming the presence of within subject variability. Also, no two subjects had identical behavior, confirming the presence of between subject variability (Fig. 1b). Therefore, our motor task successfully produced behavior that varied from trial-to-trial both within and between subjects.

**Figure 1.**
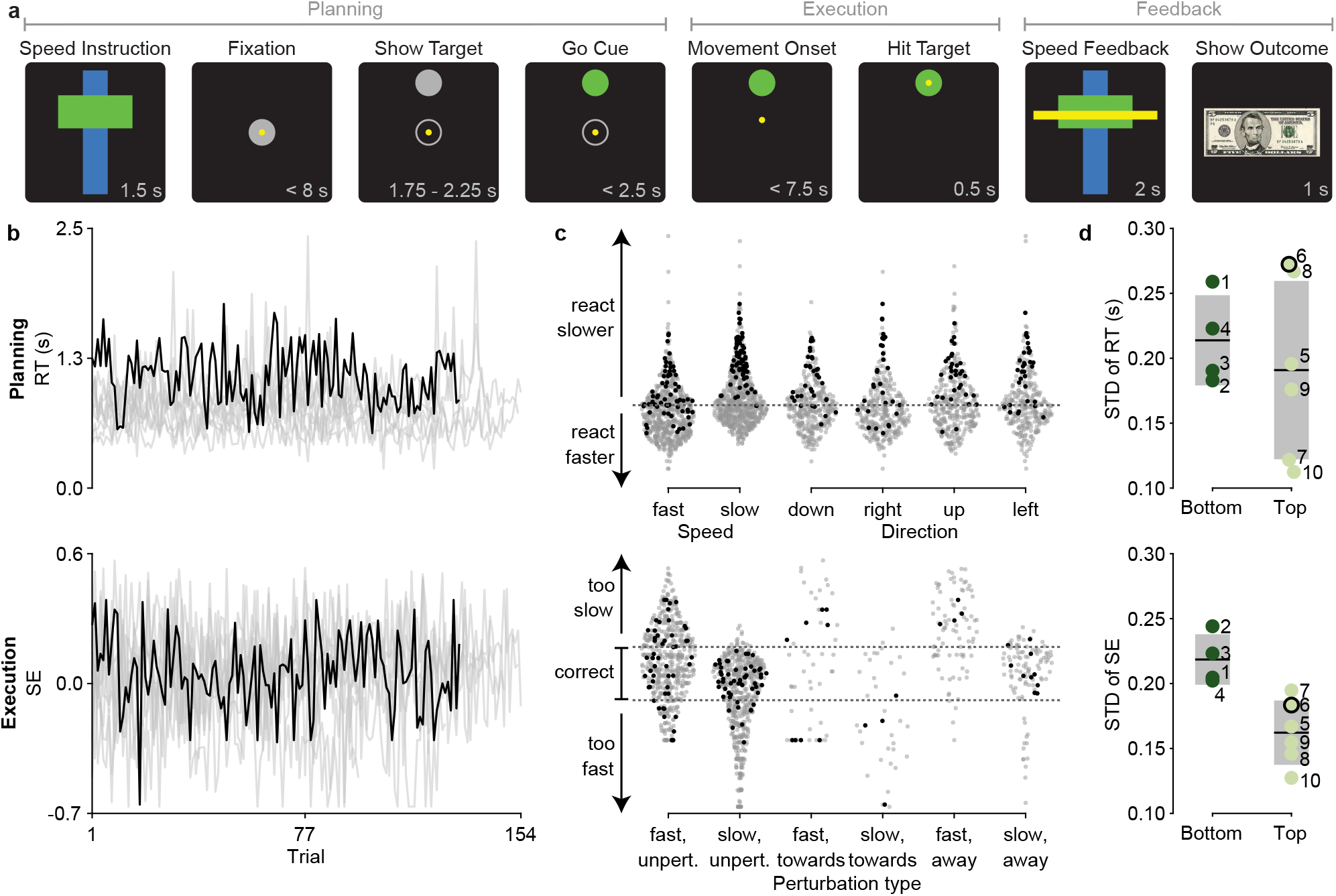
Movement variability across subjects and trial conditions for motor task. Summary of behavior during motor task shows variability within and between subjects that is independent of trial conditions yet correlates with their session performance. **a,** A detailed timeline of epochs during a single simulated trial shown using. The name of each epoch is above each visual stimuli. The interval of time in which the stimuli were presented is in the bottom right corner of each simulated screen. The trial conditions (speed, direction, perturbation) for this example are fast, up, and unperturbed. This trial is also correct. **b,** Time-series of the observed RT (top) and SE (bottom) across all trials and subjects. Subject 6 is traced in black. The remainder of the subjects are traced in gray. **c,** Observed RT (top) and SE (bottom) for all trials and subjects separated by the trial conditions. Each marker represents the behavior of a subject during a trial with the specified trial conditions. Subject 6 is colored as black. The remainder of the subjects are colored as gray. The dark gray dotted line on top plot indicates the average population RT (0.80 s). The dark gray dotted lines on bottom plot indicates the tolerance of SE between −0.13 and 0.13, where markers between these lines represents correct trials. The arrows indicate the interpretation of the behavior relative to average. **d,** Comparison of the Standard Deviation (STD) of the observed RT (top) and SE (bottom) and performance group of each subject. Each marker is labeled by the subject it represents and colored by the performance group they belong too. Subjects with session performance below average session performance of the population (51 %) are called “bottom performers” (dark green) and those below are called “top performers” (light green). Subject 6 is outlined in black. The group mean is represented by the black solid line and ±1 STD is represented by the grey box. There is a a relationship between behavior variability and session performance in which better performers are less variable.

We then investigated whether differences in trial conditions could explain the variability observed within subjects. Figure 1c shows the distributions of the RT (top) and SE (bottom) for the population separated based on the trial condition. Starting with the planning phase, the trial conditions that influenced RT were speed and direction. We expected subjects to change their RT based on these conditions speed. Indeed, subjects reacted more quickly for fast trials than slow trials (*p* = 0.014, ANOVA). They also reacted more slowly for trials with an upward motion to the target (*p* = 0.013, ANOVA). For the execution phase, the trial conditions that influenced SE were speed and perturbation. Both conditions significantly influenced the SE; subjects would move too slow for fast trials (0.00017, ANOVA) or when perturbed away (0.014, ANOVA). We also found significant differences between subject’s RT (*p* = 0.0054, ANOVA) and SE (*p* = 0.022, ANOVA). Taken together, these results support a model of planning and execution using RT and SE that is both subject-specific and based on trial conditions. Supplementary Table 4 contains the complete break down of the trial conditions each subject experienced.

We also found that performance differentiates variability between subjects. Variability was quantified using the Standard Deviation (STD) of each behavior, where a higher STD corresponds to higher variable behavior. Subjects were divided into groups based on their session performance. Subjects whose session performance was above average (51 %) as “top performers” and those below as “bottom performers”. Figure 1d shows how subject’s variability of RT (top) and SE (bottom) depends on their performance. Specifically, we found that top performers have less variable behavior.

In summary, even though all the subjects encountered the same trial conditions, their behavior varied which, in turn, affected their performance. Therefore, factors other than trial conditions must be influencing their performance.

### Internal states capture movement variability that trial conditions cannot

Using computational methods, we next developed a model to account for the variability that we observed between subjects (see Methods). Specifically, to account for variability not captured by trial conditions, we added two internal states. The first internal state was the “error state,” which accumulates the speed errors during past trials to keep track of how well a subject was accomplishing the instructed speed. The second internal state was the “perturbed state,” which accumulates the presence of perturbations during past trials to convey environmental uncertainty.

To combine trial conditions and internal states, we used the state-space model illustrated in Figure 2a. Each behavior (equations (1) and (2)) was calculated as a linear combination of the trial conditions and internal states (equations (3) and (4)). These equations were then used to simultaneously estimate the behavior and internal states for all subjects. Here, we used subject 6 to demonstrate the intuition behind our model that can be applied to the population. Specifically, we examined (i) if our estimates follows the observed behavior, (ii) the characteristics that internal states and trial conditions independently capture, and (iii) how their internal states uniquely evolve. The model results for all subjects are in Supplementary Figure 1 through Supplementary Figure 5.

**Figure 2.**
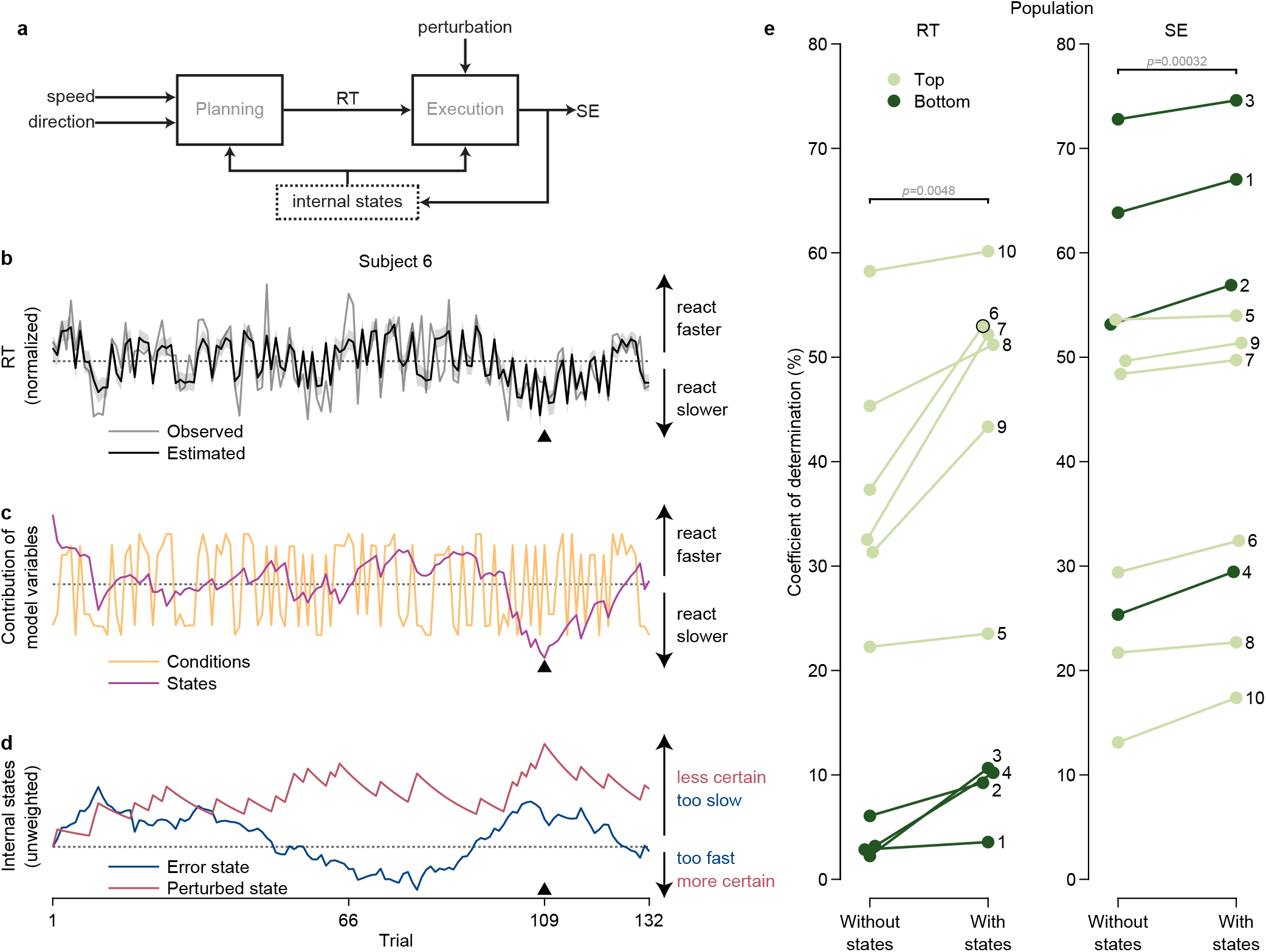
Influence of model variables for estimating behavioral model. **a,** Block diagram for our dynamical model, representing the brain, that models behavior to capture movement variability. Internal states are outlined by a dotted line to highlight its latent feedback structure in the model. Examination of the model variables for subject 6 reveals underlying latent dynamics from the internal states that leads to the improvement of estimated behavior across all subject. The triangle in panels **b-d** marks trial 109. **b,** Time-series of the observed (gray solid line) and estimated (black solid line) RT over all trials for subject 6. The shaded area around the estimated line represents the 95 % confidence bounds. The dark gray dotted line markers their average RT. **c,** Time-series of the conditions (orange solid line) and states (purple solid line) over all trials for subject 6. The dark gray dotted line markers their average RT. Adding the conditions and states together yields the estimated RT. **d,** Time-series of the error state (blue solid line) and perturbed state (red solid line) over all trials for subject 6. For demonstrative purposes, the states are not weighted but are scaled by their standard deviation. The dark gray dotted line is marked at 0. **e,** Goodness-of-fit for the RT (left) and SE (right) models across all subjects. This was measured using the coefficient of determination between the observed and estimated behavior, which ranges between 0 % (worst) and 100 % (best). We compared the behavioral models with internal states (“With states”) to a models with the same trial conditions but without internal states (“Without states”). Each marker is labeled by the subject it represents and color-coded based on their performance group (dark green for bottom performers and light green for top performers). Subject 6 is outlined in black. Adding internal states significantly improved the goodness-of-fit of the model for all subjects (*p*-values inset, paired *t*-test), as larger values are better. Top performers benefited the most from adding internal states to RT.

Overall, we first found that the estimated behavior follows the observed behavior. Figure 2b shows the RT across all trials for subject 6, with the observed in gray and the estimated in black. Visually, the estimate follows key features, including sudden jumps between trials and gradual changes such as between trials 100 and 125. For example, the estimate on trial 109 (black triangle) matches what was observed, which was that subject 6 reacted faster than average.

Next, we explored which parts of our model were responsible for capturing these features. Figure 2c shows the trial conditions (orange) and internal states (purple) used to estimate RT for subject 6. We observed that the conditions accounted for the sudden jumps between trials and states accounted for the gradual changes. For example, on trial 109 (black triangle), the conditions were slow and up, which typically caused them to react slower than average. However, the states outweighed the conditions, leading them to react faster than average. By incorporating both trial conditions and internal states, our model captured features essential in realistic behavior that neither would be able to convey independently.

To determine why subject 6 reacted faster than average despite trial conditions, we next examined the structure of the two internal states to understand their dynamics. Figure 2d shows the error state (blue) and perturbed state (red) across all trials for subject 6. The error and perturbed state on trial 109 (black triangle) are both positive, indicating that they recently moved slower than instructed and were perturbed. Hence, the internal states grant access to latent information about subjects. The perturbed state conveyed that recent successions of perturbations caused subject 6 to react and move faster in an attempt to counteract the uncertain environment.

Finally, we looked at our population of subjects to test whether adding internal states improved our ability to explain movement variability over using trial conditions alone, by comparing the coefficient of determination—a goodness-of-fit metric that measures the proportion of the behavioral variability that can be explained by the model variables—of the model with states to one without states. Figure 2e shows that adding the internal states to the model significantly improved the estimation of both RT and SE across all subjects (RT: *p* = 0.0048, SE: *p* = 0.00032, paired *t*-test), where the higher percentage means more of the variability is accounted for by the model variables. Recall that subjects were classified as either bottom (dark green) or top (light green) performers based on their session performance. In this context, it was interesting that top performers benefited the most from adding internal states to estimate RT. For example, the goodness-of-fit for subject 6 improved by 16 %. In summary, internal states are essential for completely capturing movement variability. They conveyed slow evolving characteristics in the behavior, caused by retained trial history, that were not accounted for by the trial conditions.

### Top performers learn from previous trials

The amount a subject weighs model variables reveals what they prioritized when deciding how to vary their behavior. Thus, we wondered whether top and bottom performers might use different strategies that could be uncovered using our computational approach. Specifically, we hypothesized that top performers improved their performance by learning selective information from previous trials. Indeed, we found that they learned from the error state but not the perturbed state. We tested this by considering the relationship between subjects’ weights on the internal states and session performance. Comparisons across all model variables are in Supplementary Figure 6.

Top performers reduced their RT (Fig. 3a) and SE (Fig. 3b) through the negative weights on the error state after accumulating positive errors by moving slower than instructed. Under the same circumstances, bottom performers continued to react and move more slowly than instructed due to the positive weights on the error state, accumulating more errors. These results indicate that top performers adjusted their behavior to improve while bottom performers maintained their error tendencies. This finding shows that top performers learn based on feedback.

**Figure 3.**
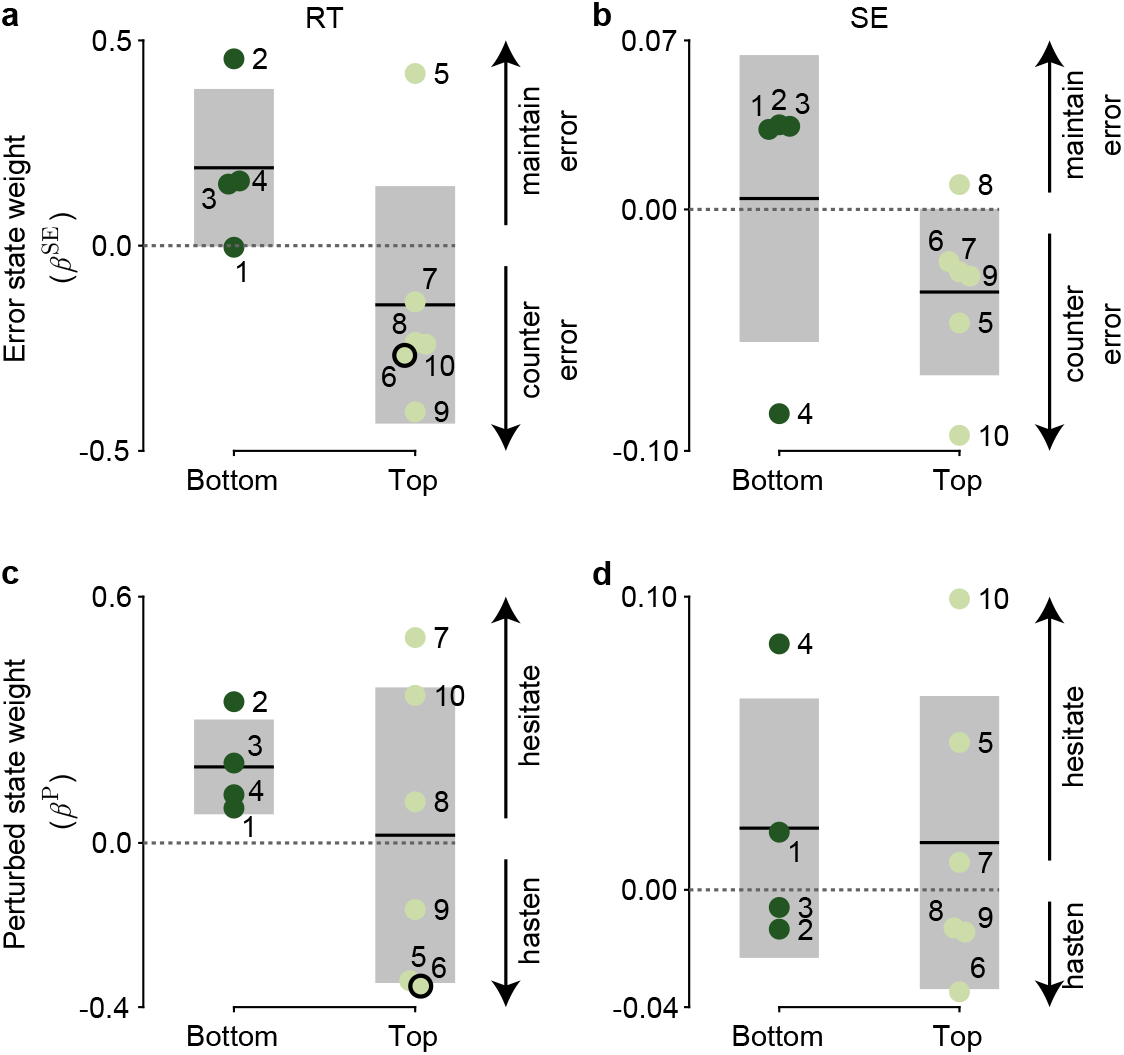
Weights on internal states show subjects use different motor strategy. Comparison between the internal states weights and performance group reveals that top performers learn from error and became more vigilant after perturbations. The larger the magnitude of the weight, the larger the impact the internal state has on behavior. The sign determines the impact the internal state has on behavior. Each marker is labeled by the subject it represents and color-coded based on their performance group (dark green for bottom performers and light green for top performers). Subject 6 is outlined in black. The group mean is represented by the black solid line and ±1 STD is represented by the grey box. The dark gray dotted lines marks where weights equal 0 (*i.e.*, internal state does not impact behavior). **a,** Top performers countered their error by reacting faster than average (negative weight) and bottom performers maintained their error by reacting slower than average (positive weight) after moving too slow (positive error state). **b,** Top performers tended to move too fast (negative weight) and bottom performers tend to move too slow (positive weight) after moving too slow (positive error state). **c,** Half of top performers and all bottom performers hesitated after perturbation trials (positive perturbed state) by reacting slower (positive weight). **d,** One half of subjects hesitated by moving too slow (positive weight) and the other half hastened by moving too fast (negative weight) after perturbations trials (positive perturbed state).

Alternatively, we did not foresee top subjects learning from perturbed trials due to their unpredictability. Indeed, Figure 3c and Figure 3d show that there was no relationship between top and bottom performers with the weight of the perturbed state on RT and SE, respectively. Instead, we found that subjects responded by either hesitating (*i.e.*, react slower or move too slow) or counteracting (*i.e.*, react faster or move too fast) for subsequent trials when they perceived the environment to be uncertain. All bottom performers and half of the top performers positively weighted the perturbed state on RT, indicating hesitation to react. Conversely, half of the top performers hastened by reacting faster, heightening their senses in preparation for a possible perturbation. In terms of SE, we found an equal mixture of performers who moved slower (positive weight) and faster (negative weight) after perturbations. We suggest that those who moved faster did so because they were exerting more force in their movement in case they were perturbed.

In summary, our model could account for differences in the strategies of top and bottom performers, where top performers learned to counteract errors directly based on previous feedback. Though we could not find a complementary strategy against perturbations, we did find that subjects either hesitated or hastened to react when they perceived the environment to be uncertain.

### Internal states are encoded by large-scale brain networks which correlate with session performance

Our combined experimental and modeling results provide support for the proposal that internal states can account for behavioral variability, within and across subjects. We next asked whether it was possible to gain insight into neural correlates of these internal states. More specifically, we asked whether such states are encoded by large-scale brain networks. To do this, we first assessed whether we could identify a set of brain regions linked to each internal state, and then determined which regions preferentially map to distinct large-scale brain networks related to session performance.

As a result of our unique experimental procedure, we had access to neural recordings from intracranial depth electrodes from each human subject simultaneously as they performed the motor task used to derive their internal states. Subjects were implanted with electrodes by clinicians to localize the epileptogenic zone for treatment. Illustrated in Figure 4a, this granted us access to local field potential activity from a broad coverage of nonmotor regions, where we hypothesized the brain encoded internal states. To identify any such region with these neural correlates, we used a non-parametric cluster statistic^[35]^ between the spectral data of each region and each internal state across the population (see Methods). This unsupervised method provided windows of time (during a trial) and frequency (between 1 and 200 Hz) in which the power of each region correlated with the internal state across trials (Figure 4b). These regions were first labeled anatomically using semi-automated electrode localization before being mapped to large-scale brain networks (see Methods).

**Figure 4.**
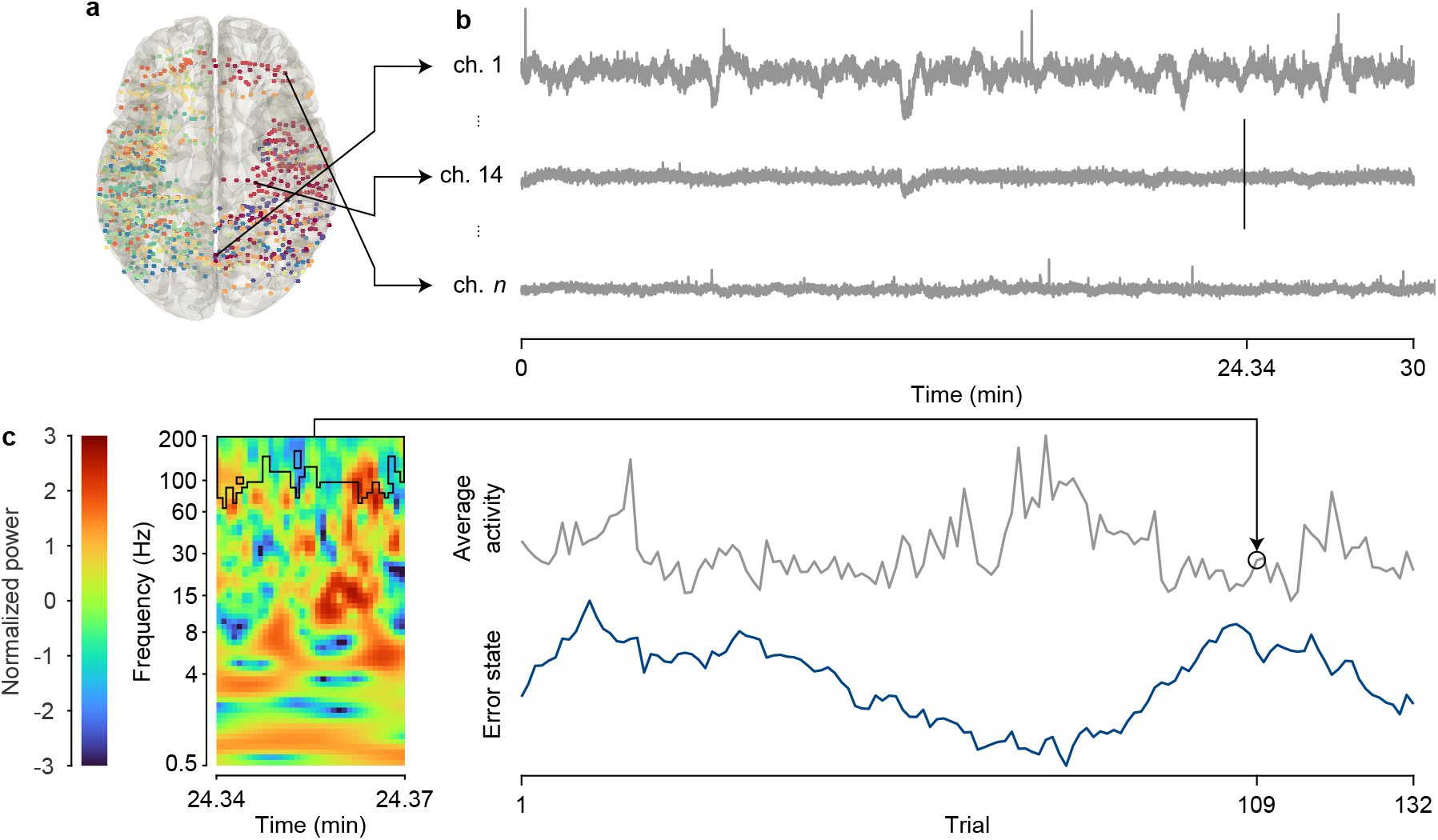
Identifying correlates between neural data and internal states. **a,** Each subject was implanted with multiple intercranial depth electrodes (SEEG). This electrode coverage can be visualized by mapping subject-specific coordinates of each channel onto a template brain. Each marker represents the coordinates of a channel from the native space of the subject warped to a common atlas space and are color-coded for each subject. Left (L) and Right (R) hemisphere are labeled. **b,** We simultaneously recorded Local Field Potential (LFP) activity (in mV) at millisecond resolution from each of these channels along the electrode, which mainly covered temporal, parietal, and limbic brain regions. The LFP activity from subject 6 is shown, with the arrows indicating the location of the channel they are recorded from. **c,** Graphical representation of non-parametric cluster statistic between neural data and internal states. Following subject 6 as an example, we found a relationship between the brain region intraparietal sulcus right (IPS R) and their error state. The spectral data (left) is represented by the time–frequency domain where the color of each pixel represents the normalized power at the specified time and frequency bin. The statistic finds time-frequency windows (left, outlined in black) in which the average power of each pixel within the window (top right) correlates with the internal states (bottom right) across trials. For example, the spectral data belongs to trial 109 (marked by the vertical black line in **b** at 24.34 min) and the average normalized power in the outlined window is circled (top right).

To determine which regions encode each internal state, we chose those whose power significantly correlated with the state from trial-to-trial in the population. Our modeling results showed that subjects weighed internal states differently, primarily based on their session performance. We suspected that this would be reflected in the brain by how well these regions encoded the states. The degree to which a subject encodes an internal state in a region, which we called the encoding strength, was quantified by correlating the average power within the time-frequency window (from the population statistic) to the state on a trial-by-trial basis (see Methods). We used this to isolate a set of regions whose activity modulated with the state and whose encoding strength correlated with session performance.

#### Error state is encoded by the dorsal attention network

First, we found that the error state was primarily encoded by regions in the DAN (see Table 2 for details and Fig. 5a for visualization). The regions in DAN (blue) included the intraparietal sulcus right (IPS R), middle temporal gyrus right (MTG R), superior frontal gyrus left (SFG L), superior parietal lobule left (SPL L), and superior temporal sulcus right (STS R). The second most prominent network was the visual network (yellow), which included the parieto-occipital sulcus right (POS R), anterior transverse collateral sulcus left (ATCS L), cuneus right (Cu R), and inferior temporal gyrus right (ITG R). A couple of regions from other networks also appeared, including the supramarginal gyrus right (SMG R) from the Ventral Attention Network (VAN) as well as some from the DN.

**Figure 5.**
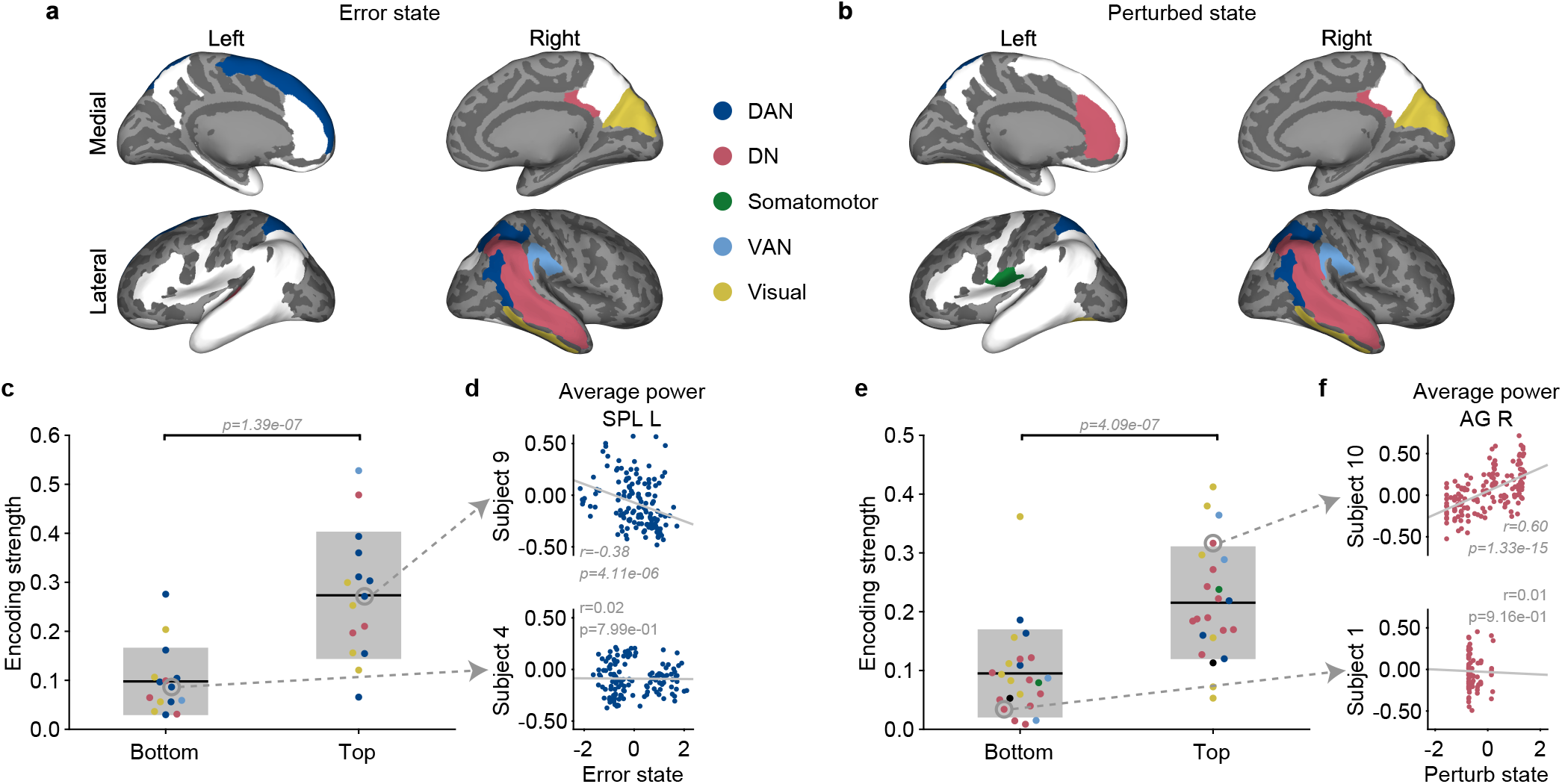
Neural activity in large-scale brain networks that encode internal states scales based on session performance. Summary of regions found by our analysis that encode the **a,** error state and **b,** perturbed state. They are highlighted on an inflated template brain based on the large-scale brain network they belong to: DAN in dark blue, DN in red, Somatomotor in green, VAN in light blue, and visual in yellow. The light gray areas represent the gyri and the dark gray areas represent the sulci. The white areas represent regions included in our analysis but did not encode the state. Top and bottom performers encoded the **c,** error state and **e,** perturbed state to different degrees. Each marker represents the average encoding strength in a region that encodes the state for each performance group and is color-coded by the network the region belongs to. An encoding strength of 1 means the state is strongly encoded by the activity of a region whereas 0 means the state is not encoded. The average encoding strength of the performance group is represented by the black solid line and ±1 STD is represented by the grey box. We found that top performers had neural activity that modulated significantly more with either internal states than bottom performers (*p*-value in panel, two-sample *t*-test). Examples of data used to calculate the encoding strength of a top (top) and bottom (bottom) performer for the **d,** error state and **f,** perturbed state. The left superior parietal lobule (SPL L) between top subject 9 and bottom subject 4 in the DAN represents the error state and the right angular gyrus (AG R) between top subject 10 and bottom subject 1 in the DN represents the perturbed state. Each marker represents a trial, with the corresponding neural activity of the cluster for a channel in the specified region and normalized state value. The magnitude of this correlation was used to calculate the encoding strength. The light gray solid line represents the least-square line. The Spearman correlation (*r*) and *p*-value (*p*) are included in each panel.

**Table 2.**
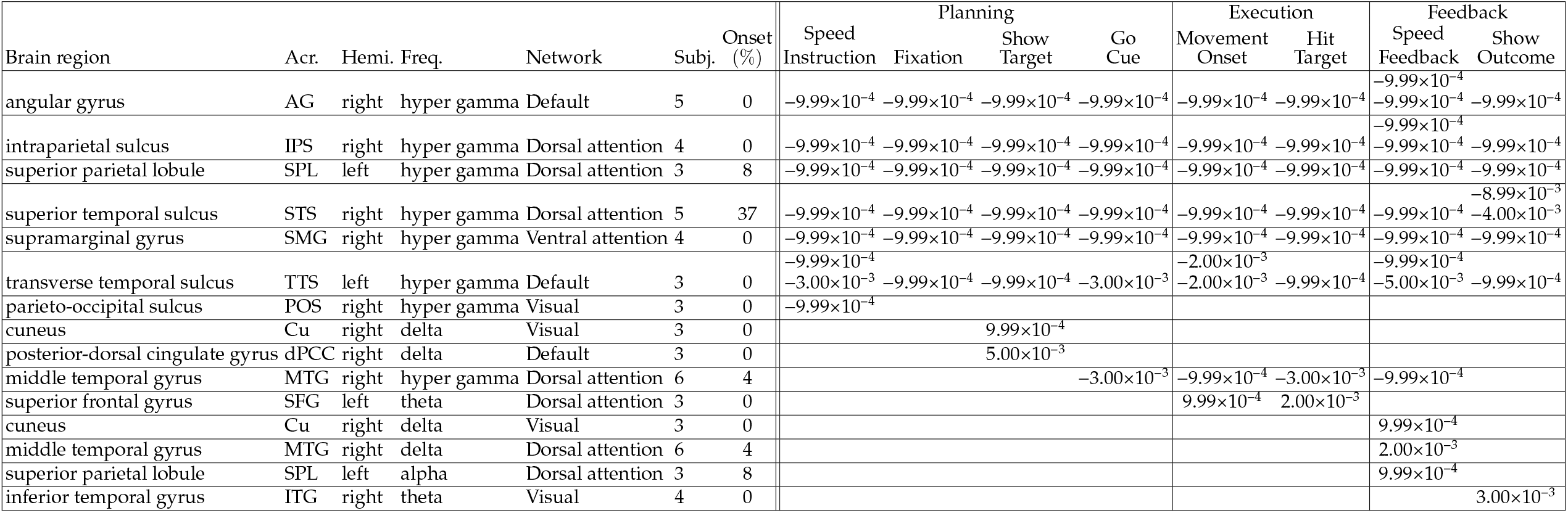
List of significant clusters across brain regions and epochs encoding the error state. For each brain region and task epoch, the table reports all clusters that pass a false discovery rate of level *q* = 0.015 and their level within the cell. For each brain region, the table also provides the acronym (Acro.), hemisphere (Hemi.), dominant frequency band (Freq.), network, number of subjects with recordings in this region (Subj.), and percentage of electrodes in this region that have been annotated as part of an onset zone (Onset).

Two groups of regions emerged based on when during the trial did their activity correlate with the error state and in what frequency band, as defined in the Methods). Half of these regions encoded the error state throughout the session as persistent activity in the frequency band hyper gamma (100–200 Hz). This activity was negatively correlated with the error state, meaning the DAN exhibited higher deactivation when a subject moved slower than instructed. The other half encoded the error state as phasic activity during either planning, execution, or feedback in frequency bands lower than 15 Hz. This activity was positively correlated with the error state for most regions. Overall, the regions in DAN use both persistent and phasic activity to encode.

Recall our result above showing that top and bottom performers used opposing strategies regarding how they used their error state to change how they reacted in future trials (Fig. 3a–b). We next investigated whether this distinction would be reflected by how strongly the DAN encodes the state. Since top performers learned from their errors, we expected a stronger correlation with neural activity as compared to the bottom performers. Indeed, consistent with our hypothesis, this trend is shown in Figure 5c, with DAN in blue. For example, the SPL L—a key hub of DAN—modulates its neural activity based on the error state for top performers (Fig. 5d(top)) but not bottom performers (Fig. 5d(bottom)). The other networks highlighted in Figure 5c show a similar trend.

#### Perturbed state is encoded by the default network

Second, we further found that the perturbed state was primarily encoded by regions in the DN (see Table 3 for details and Fig. 5b for visualization). The regions in DN (red) included the angular gyrus right (AG R), anterior cingulate gyrus left (ACG L), middle temporal gyrus right (MTG R), posterior-dorsal cingulate gyrus right (dPCC R), and superior temporal sulcus right (STS R). As with the error state, regions in the visual network (yellow) also encoded the perturbed state. They consisted of the cuneus right (Cu R), inferior temporal gyrus right (ITG R), fusiform gyrus left (FuG L), and parieto-occipital sulcus right (POS R). Other networks that appeared included the DAN (IPS R and SPL L), VAN (SMG R), and even the somatomotor network through subcentral gyrus left (SubCG L). The subcortical region hippocampus right (Hippo R) briefly encoded the perturbed state after execution. However, since we were only interested in large-scale brain networks of the neocortex, Hippo R was not classified in this study.

**Table 3.**
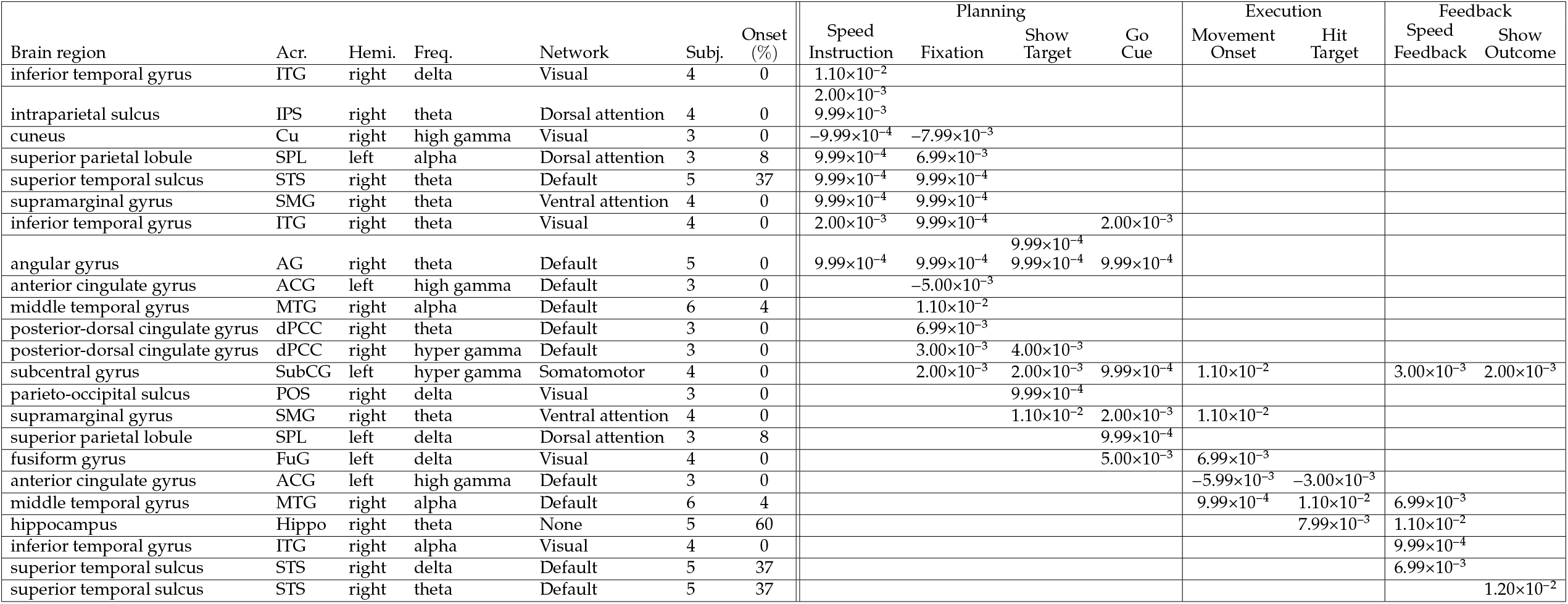
List of significant clusters across brain regions and epochs encoding the perturbed state. For each brain region and task epoch, the table reports all clusters that pass a false discovery rate of level *q* = 0.015 and their level within the cell. For each brain region, the table also provides the acronym (Acr.), hemisphere (Hemi.), dominant frequency band (Freq.), network, number of subjects with recordings in this region (Subj.), and the percentage of electrodes in this region that have been annotated as part of an onset zone (Onset).

The regions in DN that encoded the perturbed state did so with phasic activity—activity during planning, execution, or feedback—in frequency bands under 15 Hz (*i.e.*, delta, theta, alpha). Most regions had activity that was positively correlated with the perturbed state; trials with a high perturbed state (associated with recent perturbations) coincided with higher activation of DN.

Recall our earlier result in which subjects alter their behavior (*i.e.*, hesitate or hasten) with regards to the perturbed state. Although there was no consistent strategy between performers, we still speculated whether there would be a neurological difference between performers in how the perturbed state was encoded. As with the error state, we expected top performers to encode the perturbed state more because they generally weighted the perturbed state with a higher magnitude than bottom performers (Fig. 3c–d). Indeed, Figure 5e shows that all networks, namely the DN (red), modulate their neural activity based on the perturbed state for top performers but not for bottom performers. For example, as a hub for the DN, the AG R would increase activity when the perturbed state was high (*i.e.*, after perturbation trials) for top performers (Fig. 5f)(top)) but not for bottom performers (Fig. 5f)(bottom)). Even though our models could not find a consistent strategy between performers, top performers still modulated their activity to match the perturbed state.

### Connectivity strength within networks correlates to motor performance and strategy

Recall that we showed above that a subject’s weight on their internal states provides insight into their learning strategies. Since subjects encode the internal states in distinct networks, we hypothesized that these strategies would further be reflected by the functional connectivity of the networks that we identified as encoding both the internal state and performance. Simply put, pairs of regions that are spatially separate yet whose neural activity is correlated are functionally connected^[36]^. Like encoding strength, we quantified this relationship between pairs, which we called the connectivity strength, by correlating their average power within the time-frequency window (from the population statistic) to each other on a trial-by-trial basis (see Methods). Unfortunately, we were not able to observe all possible pairs of regions because either the pair was not represented in the data set, or we did not have enough subjects with the pair (*n ≥* 3) to make any remarks. Since connectivity strength is also subject-specific, we expected top and bottom performers to use different connections to implement different strategies. Since we found that they encoded the internal states stronger than bottom performers, we focused on connectivity favored by top performers.

Our results above demonstrated that top performers improved their performance by compensating their behavior in response to the error state. Using the set of regions that encode the error state, Figure 6a further shows that top performers favored connectivity between regions in the DAN (blue), VAN (light blue), and visual network (yellow). We found that regions that encoded with persistent activity connected to regions that encoded with phasic activity. Specifically, the persistent activity from DAN (IPS R and SPL L) and VAN (SMG R) projected namely to regions in the visual network during key phases during the trial (*i.e.*, planning, execution, feedback). This result suggests that the error state is held and distributed by the attention networks to modulate visual attention. An increase in such attention could account for the counteracting behavior observed by top performers, such as reacting faster after moving slower than instructed (Fig. 3a). We also observed a significant correlation between the persistent activity and phasic activity during feedback for the SPL L, which is a key hub of DAN. This relationship implicates persistent and phasic activity having different roles when encoding the error state. Perhaps the phasic activity represents the various sensory processes (depending on the phase) used to extrapolate information about the error state and the persistent activity holds this information in memory for accessibility by other regions. Then, the connection between SPL L could be an example of updating between integration and memory. Overall, top performers favored connections between DAN and visual networks to encode the error state. These connections could account for how top performers learned from their errors, specifically by updating their memory based on visual feedback to modulate attention in future trials.

**Figure 6.**
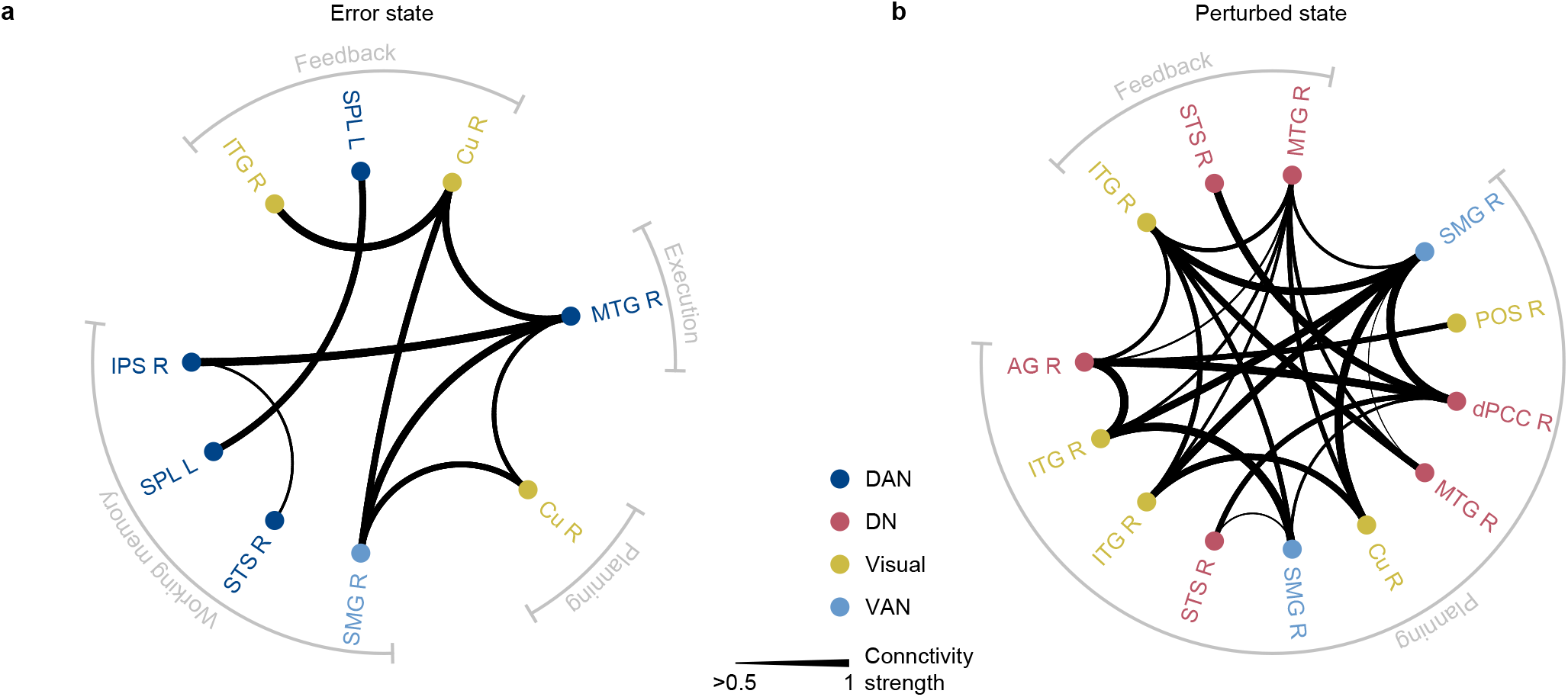
Network connectivity of top performers supports learning strategy. Directed graph depicting the strength of the relationship between connectivity strength and session performance for each pair of regions. The regions are color-coded by the large-scale brain network they belong to: DAN in dark blue, DN in red, VAN in light blue, and visual in yellow. Only the pairs of regions whose connectivity strength related to session performance are shown. The magnitude of the correlation between connectivity strength and session performance are depicted by the thickness of the edge; the higher the correlation, the thicker the edge with values truncated after 0.5. The regions are ordered by when they encode each state and its phase is labeled along the circumference, demonstrating synchrony across time. **a,** Functional connectivity for top performers between pairs of regions that encode the error state. **b,** Functional connectivity for top performers between pairs of regions that encode the perturbed state.

Our results above also demonstrated that top performers altered how they react (*i.e.*, hesitate or hasten) in response to the perturbed state. Figure 6b further shows the connectivity between the perturbed state encoding regions favored by top performers. Notable, we found more connections for the perturbed state than compared to the error state. This makes sense because the perturbed state was more disruptive to their behavior and required the integration of sensorimotor pathways for interpretation during planning and feedback. In general, most of the connections we found were between DN (red) and visual network (yellow). This relationship could allow for the communication of visual information from feedback to be available for the DN during planning, such as to update their expectation about the environment of the motor task. This information could then be projected to regions in the visual network and VAN to plan their future behavior. For example, dPCC R (hub of DN) during early planning projects to SMG R (hub of VAN) during late planning and early execution. Taken together, these results suggest that the DN could be modulating bottom-up visual attention based on the perturbed state. Overall, top performers favored connectivity between DN to other relevant networks whose activity modulates based on the perturbed state during planning and feedback, which they could have used to adapt their responsiveness based on perturbations we observed from their behavior.

## Discussion

In the present study, we first identified two internal states—based on error and environmental history—that induce movement variability in humans. The degree to which states contribute to an individual’s variability reveals opposing strategies between the top and bottom performers regarding how they used their states to inform future behavior. Remarkably, we then found that these internal states were linked to encoding in large-scale brain networks, DAN and DN, respectively. Taken together our findings reveal that differences in large-scale brain networks that can distinguish top from bottom performers: (i) top performers modulate network activity on a trial-by-trial basis with respect to their internal states and (ii) their learning strategy is supported by explicit connections within and between networks during phases of movement.

### Internal states and variability in motor control

The general effect of error and the environment on movement variability are well documented in motor control. Traditionally, the goal of optimal feedback control is to minimize error during movement, where larger errors require more variability to correct the movement^[37, 38]^. By accounting for the accumulation of errors from trial-to-trial, we also found that variability in top performers scaled based on error history. In everyday life, we adapt our behavior to fit the environment based on prior experience. But it is difficult to adapt when disturbances are rare and unpredictable. Fine and Thoroughman found that it is difficult for subjects to learn how to respond to these disturbances that occur for less than 20% of trials^[39]^. They proposed an adaptive switch strategy that depends on the environmental dynamics: ignore performance from trials with rare disturbances and learn when they are common^[40]^. Complimentary, we found that subjects responded in trials after perturbations by either reacting hesitantly or vigorously. Nevertheless, their strategy was not a predictor of how well they performed our reaching task, which can be further explored in future studies.

### Modulation of dorsal attention network with error history and links to performance

To learn our speed-instructed motor task, subjects were required to keep track of past errors using working memory. We found that our subjects learned the task by monitoring their history of errors across trials to decide where to allocate attention through the DAN. Specifically, we found DAN activated in frequencies below 15 Hz and deactivated in frequencies above 100 Hz when subjects recently moved faster than instructed to accumulate positive error, to which they would then slow down. These findings are in line with the speed-accuracy trade-off phenomenon which is observed during motor learning in the form of behavioral variability where performance is optimized by balancing moving faster at the cost of making more errors^[41]^ which innately requires tracking history^[42]^. For the first time, our results implicate DAN as a network as encoding tracking history. Increases in Blood-Oxygen-Level-Dependent (BOLD) signal relative to baseline in DAN have been reported when subjects were instructed to prioritize speed (instructed as fast or slow) over accuracy during a response interference speed-accuracy trade-off task^[43]^. Another study using an anti-saccade task found DAN activation through BOLD signal positively correlated with RT (*i.e.*, more activation when slowing down) and being the least activated on trials with large errors (which compares to our task when the error state would be close to zero)^[44]^. In a visually guided motor sequence learning task, DAN activated—through BOLD signal—to large errors during early learning, which they related to active visuospatial attention when first learning the sequence^[45]^.

We also found that the DAN encoded error history throughout the trial using persistent activity in frequencies above 100 Hz. Such activity is characteristic of working memory^[46–48]^. During motor planning in tasks with working memory, DAN has been shown to maintain task-relevant information, such as target location, during delay periods using persistent activity in both whole-brain and single unit recordings^[49–51]^. DAN has also been linked to working memory closely tied to visual attenuation related to memory load and top-down memory attention control during visual working memory tasks^[52]^. Though our task does not explicitly study working memory, given the evidence, our results suggest that DAN is tracking the accumulation of past errors in working memory. This would also support our connectivity results as information held in working memory can be easily accessed by multiple systems, such as for sensorimotor integration or visual processing, for recalling and updating^[53–55]^.

Finally, we found that the encoding strength of regions in the DAN and functional connectivity between these regions scaled based on the subject’s overall performance. We observed that top and bottom performers have different strategies. Top performers more strongly encoded error history in and between the regions in the DAN. Hence, they were more engaged in the task and modulated their attention based on the error state. This led to them slowing down after they moved too quickly and vice versa as predicted by their model weights. Meanwhile, bottom performers have poor attentional control and memory capacity and thus did not learn from their mistakes. Studies have shown that poor overall behavioral performance is related to decreased activity in DAN^[56–60]^ (called “out of the zone” ^[61]^), fluctuations in attention and working memory known as “lapse in attention”^[62–64]^, and poor connectivity in DAN^[65, 66]^. Clinically, studies of Attention Deficit Hyperactivity Disorder (ADHD) found compromised performance during working memory tasks is related to poor attentional control in DAN-related regions^[67, 68]^, similar to what we observed in our poor performers.

### Modulation of default network with environmental uncertainty and links to performance

The random perturbations made our task more difficult by creating uncertainty in the environment. Our models show that subjects also kept track of past perturbations suggesting possible attempts to learn the environment. We found that the regions whose activity correlated with the perturbed state were in the DN and did so primarily in the frequency bands theta and alpha (below 15 Hz). That is, regions in the DN encoded the perturbed state by increasing activity when the environment was perceived to be more uncertain. This function of the DN is similar to a recent study by Brandman et al. in which they found that the DN activated immediately following unexpected stimuli in the form of surprising events during movie clips^[69]^. They suggested that the DN could be involved in prediction-error representation, which our results also support. Furthermore, these authors found that DN also coactivated with the hippocampus during unexpected stimuli. This finding parallels a previous report from our dataset which demonstrated activation of the hippocampus activated in response to motor uncertainty^[70]^. A proposed process model of reinforcement learning incorporates regions in the DN and hippocampus that predict and evaluate the semantic knowledge about the environment to inform future behavior^[71]^. Taken together, our findings suggest that we captured the DN responding to the unexpected stimuli by updating semantic knowledge about the environment which informs future behavior based on the perturbed state.

Behaviorally, our model did not establish a link between subjects’ performance and how they handled past perturbations but our analysis of neural activity revealed that top performers demonstrated increased activation of regions in DN in response to the history of perturbations as well as correlated activity across regions between trials. Since subjects have no control over the perturbations, we speculate that the only thing they can do is to learn how to react in a way that optimizes their performance. Although they are applied to random trials and in random directions, perturbations are always applied during the beginning of the movement. Therefore, subjects can learn to prepare themselves in a way they see fit. Our findings suggest that top performers effectively implement their new semantic knowledge about the environment to explore different approaches to prepare for the possibility of future perturbations. DN becomes more activated during early learning^[45]^, particularly when motor imaginary is used^[72]^. In fact, athletes (*i.e.*, experts or top performers) have been shown to activate the DN when employing strategies that decrease variability, resulting in stable performance during movement^[66]^. The phenomenon, known as “in the zone”^[61]^, has been linked to the DN activation with consistent performance associated with preparedness^[66,73,74]^ and vigilance^[75]^. Hence, activation of DN indicates those subjects are prepared for the chance of a perturbation. Taken together, these findings suggest that top performers react to uncertainty by heightening vigilance through activation and connectivity in DN.

### General implications for understanding motor control

Observing movement variability in the form of motor error is common in motor control, with paradigms typically focused on aspects of motor learning. Numerous motor control studies have found—directly or indirectly—the involvement of regions in DAN and working memory^[50,54,76]^. In fact, our results align with classic motor control reports when considering their results in terms of networks. For example, Diedrichsen et al. identified neural correlates of error in DAN and visual networks, represented by SPL and POS respectively^[77]^, identical to ours. Gnadt & Anderson observed persistent activity in IPS (hub for DAN) in relation to target location during the delay in motor planning, connecting their results to memory^[49]^. In a study directed towards large-scale brain networks, DAN activation during early learning was correlated with decreased error rate, which they related to active visuospatial attention when first learning the sequence^[45]^.

### Study limitations

This study highlights the complexity of behavioral and neural data have as well as how challenging it is to disentangle internal states from other processes, nevertheless, we acknowledge several limitations inherent to our approach. First, it is possible that some of our neural data that includes results from epileptic brain regions in which activity could differ from comparable regions in healthy humans, despite precautions we took to minimize this possibility as discussed in Methods. The effects of anti-epileptic medications is another confounder that can influence the magnitude of the results, though subjects ceased their medications during clinical investigation. At this time, the only ethical method to record from the brain necessary for our study using SEEG depth electrodes in humans is while they are implanted for clinical purposes. Second, our behavioral data was limited by the design of the motor task and trial conditions, including two speeds and four directions. Future experiments could explore other trial conditions, like introducing obstacles into the task space or other stimuli such as audio. One could also imagine designing a study that picks trial conditions to produce desired variability from a subject based on model inferences about their internal states. Third, we focused on using a simple modeling approach, which raises the possibility that a key factor in a subject’s behavioral variability may be absent from our model. Contrarily, this simplicity allows for other variables, such as other trial conditions or internal states, to be easily designed and integrated to create a model for a variety of behavioral tasks.

## Conclusion

In conclusion, our findings provide a new viewpoint for motor control research. Our results raise the possibility that measured behavior from devices such as smartphones could be used to make inferences about a person’s brain state without needing to collect electrophysiological data, saving time and money in the health field for personalized medicine^[78]^ or business ventures such as sports^[79]^. Future studies in motor control should consider the effect of these networks on motor control and should account for the effects of internal states as we found that they play a significant role in governing behavior and its variability.

## Acknowledgements

The authors would like to thank Juan Bulacio, Jaes Jones, Hyun-Joo Park, and Susan Thompson who collected, deidentified, transferred, and anatomically labeled the neural data as well as Matthew S.D. Kerr, Kevin Kahn, and Matthew A. Johnson who designed and collected the behavior data. We also thank Amy J. Bastian and Vikram S. Chib for our discussions during thesis meetings over the years that helped propel this work forward. Finally, thank you to Jesse A. Smith for his extensive help proofreading and overarching moral support. Part of this research project was conducted using computational resources and scientific computing services at the Maryland Advanced Research Computing Center (MARCC). This work was supported by National Science Foundation grant (EFRI-MC3: #1137237).

## End Notes

### Author Contributions

M.S.B., J.A.G.-M., J.T.G., and S.V.S. designed research; M.S.B., J.A.G.-M., J.T.G., and S.V.S. performed research; M.S.B, P.S., and S.V.S. contributed analytic tools; M.S.B., P.S., and S.V.S. analyzed data; M.S.B., P.S., Z.F., J.A.G.-M., and J.T.G. provided resources and data curation; K.E.C, J.A.G.-M., and S.V.S. supervised the work; and M.S.B., P.S., Z.B.F., J.T.G, K.E.C., J.A.G.-M., and S.V.S. wrote the paper.

## Online Methods

### Recording neural data from humans

Ten human subjects (seven females and three males; mean age of 34 years) were implanted with intracranial SEEG depth electrodes and performed our motor task at the Cleveland Clinic. These subjects elected to undergo a surgical procedure for clinical treatment of their epilepsy to identify Epileptogenic Zone (EZ) for possible resection. Details of the demographic and clinical information of each subject are listed in Table 1. All experimental protocols were approved by the Cleveland Clinic Institutional Review Board. Subject criteria required volunteering individuals to be over the age of 18 with the ability to provide informed consent and able to perform the motor task. Other than the experiment, no alterations were made to their clinical care. We excluded two additional subjects who attempted to perform the task but failed to complete it.

**Table 1.**
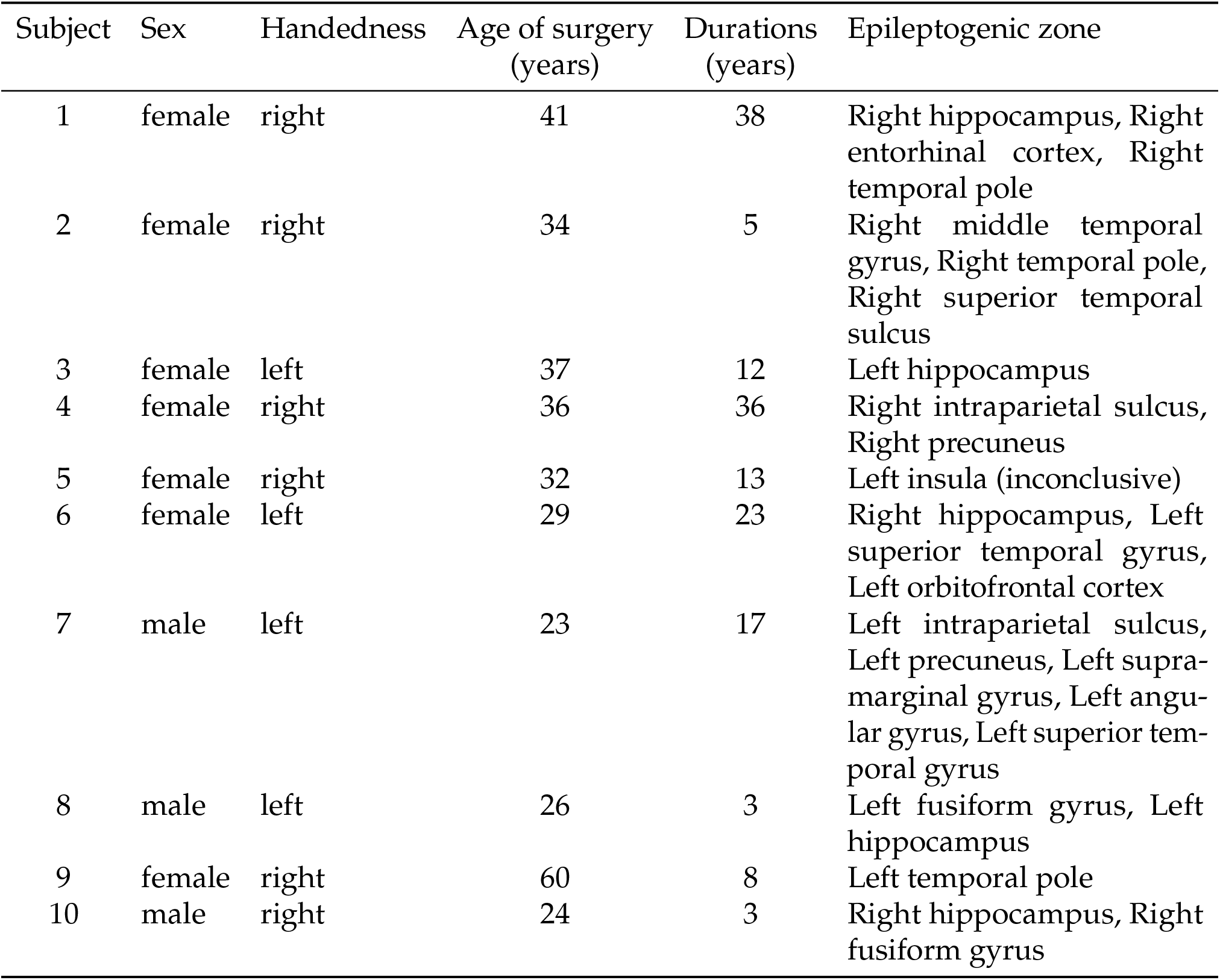
Subject demographic and clinically relevant information such as clinically identified epileptogenic zone.

| Subject | Sex | Handedness | Age of surgery<br>(years) | Durations<br>(years) | Epileptogenic zone |
| --- | --- | --- | --- | --- | --- |
| 1 | female | right | 41 | 38 | Right hippocampus, Right entorhinal cortex, Right temporal pole |
| 2 | female | right | 34 | 5 | Right middle temporal gyrus, Right temporal pole, Right superior temporal sulcus |
| 3 | female | left | 37 | 12 | Left hippocampus |
| 4 | female | right | 36 | 36 | Right intraparietal sulcus, Right precuneus |
| 5 | female | right | 32 | 13 | Left insula (inconclusive) |
| 6 | female | left | 29 | 23 | Right hippocampus, Left superior temporal gyrus, Left orbitofrontal cortex |
| 7 | male | left | 23 | 17 | Left intraparietal sulcus, Left precuneus, Left supra-marginal gyrus, Left angular gyrus, Left superior temporal gyrus |
| 8 | male | left | 26 | 3 | Left fusiform gyrus, Left hippocampus |
| 9 | female | right | 60 | 8 | Left temporal pole |
| 10 | male | right | 24 | 3 | Right hippocampus, Right fusiform gyrus |

Each subject was implanted with 8–14 stereotactically-placed depth electrodes (PMT^®^ Corporation, USA). Each electrode had between 8–16 electrode channels (henceforth referred to as *channels*) spaced 1.5 mm apart. Each channel was 2 mm long with a diameter of 0.8 mm. Depth electrodes were inserted using a robotic surgical implantation platform, (ROSA^®^, Medtech^®^, France) in either an oblique or orthogonal orientation. This procedure granted access to broad intracranial recordings in a three-dimensional arrangement, which included lateral, intermediate, and/or deep cortical as well as subcortical structures^[80]^. The day prior to surgery, volumetric preoperative Magnetic Resonance Imaging (MRI) scans (T1-weighted, contrasted with Multihance^®^, 0.1 mmol kg^−1^ of body weight) were obtained to plan safe electrodes trajectories that avoided vascular structures preoperatively. Postoperative Computed Tomography (CT) scans were obtained and coregistered with preoperative MRI scans to verify electrode placement postoperatively following implantation^[80]^. Electrophysiological data in the form of Local Field Potential (LFP) activity (Fig. 4b) were collected onsite in the Epilepsy Monitoring Unit (EMU) at the Cleveland Clinic using the clinical electrophysiology acquiring system (Neurofax EEG-1200, Nihon Kohden, USA) with a sampling rate of 2 kHz referencing an exterior channel affixed to the skull. Each recording session was also determined to be free of any ictal activity.

### Inducing movement variability using our motor task

Our motor task was a center-out delay arm reach where subjects won virtual money by controlling a cursor on a screen to reach a target with an instructed speed despite a chance of encountering a random physical perturbation^[70,81–83]^. Subjects performed this task in the EMU using a behavioral control system, which consisted of three elements: a computer screen, an InMotion2 robotic manipulandum (Interactive Motion Technologies, USA), and a Windows-based laptop computer^[81]^. The computer screen (640*×*480 px) was used to display the visual stimuli to the subject. Subjects were seated approximately 60 cm in front of the screen. The robotic manipulandum allowed for precise tracking of the arm position in a horizontal two-dimension plane relative to the subject. The subject used the robotic manipulandum to control the position of a cursor on the computer screen during the motor task restricted to a horizontal two-dimension plane relative to themselves. The laptop computer ran the motor task using a MATLAB-based software tool called MonkeyLogic^[84, 85]^.

During the session, subjects would complete as many trials as they could in 30 min. A complete trial consisted of eight epochs, each distinguishable with unique visual stimuli shown in Figure 1a. Subjects began each trial with an instructed speed, indicating whether they were supposed to move fast or slow (Speed Instruction). Next, subjects moved their cursor to a target in the center of the screen (Fixation). Once centered, subjects were presented with a target in one of the four possible directions (Show Target). A random delay was applied here in which subjects could not move their cursor out of the center until cued to do so. This cue was signaled as the target changing color from grey to green (Go Cue). After their cursor left the center (Movement Onset), there was a chance that a constant perturbation would interrupt their movement. Subjects were still expected to reach the target with the correct speed despite the perturbation. Once they reached (Hit Target) and held their cursor in the target, subjects were immediately presented with feedback of their trial speed compared to the instructed speed (Speed Feedback). The reward they were shown depended on if they matched the instructed speed or not (Show Outcome). An image of an American $5 bill was presented for correct trials while the same image overlaid by a red “X” was presented for incorrect trials. It should be noted that subject did not receive any monetary reward for participating in this task. Epochs are structured into traditional phases of motor control based on the design of the experiment. *Planning* includes Speed Instruction, Fixation, Show Target and Go Cue, *Execution* includes Movement Onset and Hit Target, and *Feedback* includes Speed Feedback and Show Outcome.

Subjects could fail a trial for any of the following reasons: not acquiring the center during Fixation, leaving the center before Go Cue, failing to leave the center after Go Cue, or inability to reach the target during Movement Onset. Regardless of the reason, the rest of the trial was aborted and subjects were presented with a red “X” before moving to the next trial.

We only used completed trials (*i.e.*, trials in which the Speed Feedback was reached) for our study. Subjects were aware that perturbations would be applied. Additionally, subjects were allowed as much time as they wanted to practice the motor task before the session began, which included the speeds, directions, and perturbations.

At the end of each session, the *session performance* of each subject was calculated as 100 ∗ (Number of completed trials with correct speed)*/*(Number of completed trials). Session performance of 0 % means the subject achieved the correct speed on none of their trials and session performance of 100 % means the subject achieved the correct speed on all of their trials. To differentiate the performance of a trial (*i.e.*, correct or incorrect) from the session (*i.e.*, percent of correct trials), we refer to the former as *trial performance*. Finally, we grouped subjects by comparing their session performance to the average session performance of the population (51 %). Those who performed higher than average are called “top performers” and those who performed lower than average are called “bottom performers”.

There are three trial conditions that varied from trial-to-trial: the instructed speed, the instructed target direction, and the type of perturbation. *Speed* refers to the binary condition categorizing the instructed speed (fast, slow). Either speed was equally likely for each trial. In actuality, the categorical representation of speed translates to a range of values based on the percent of a subject-specific maximum speed measured during calibration; when they were told to move the cursor “as quickly as possible” from the center to a right target over five trials just before starting the experiment. Fast trials accepted 66.67±13.33 % and slow trials accepted 33.33±13.33 % of their calibration speed. *Direction* refers to the four possible locations of the target relative to the center of the computer screen (down, right, up, left). The probability of each location was equally like for each trial. *Perturbation* refers to the type of perturbation, if any, that was experienced during the trial (unperturbed, towards, away). Each trial had a 20 % probability of a perturbation being applied with a random force between 2.5 to 15 N at a random angle, both selected from a uniform distribution. The perturbation was physically applied to the subject using the robotic manipulandum and would persist until the subject was shown their feedback. Perturbations can be categorized based on the angle it was applied relative to the target direction: towards or away. All other trials are unperturbed. The summary of the trial conditions experienced by each subject are listed in Supplementary Table 4.

In addition to trial conditions, we also tracked two continuous values that incurred movement variability from trial-to-trial: *Reaction Time (RT)* and *Speed Error (SE)*. The RT (in seconds) was quantified as the time it took for a subject to move their cursor out of the center after Go Cue. The SE was quantified as the difference between the middle of the range of the instructed speed (0.33 or 0.67) and their trial speed. The trial speed was found by dividing the constant distance between the center and the target (in pixels) with the total time between Go Cue and Hold Target (in seconds), then scaling it by their calibration speed. The SE can take on a value between −0.67 to 0.67, where a <u>positive</u> SE means the subject moved <u>slower</u> than instructed (*i.e.*, too slow), a <u>negative</u> SE means the subject moved <u>faster</u> than instructed (*i.e.*, too fast), and a SE between −0.13 and 0.13 means the subject was within the acceptable range for the trial to be correct. The statistics of the RT and SE for each subject are listed in Supplementary Table 5.

### Estimating internal states to capture movement variability

We sought to construct a behavioral model to capture movement variability based on data collected during our goal-directed center-out delay arm reach motor task. The behavioral data consisted of any quantifiable measurements from the motor task, namely the trial conditions, RTs, and, SEs.

Our system follows the framework outlined in Figure 2a. It takes on the structure of a state-space representation and consists of three basic elements for each trial *t*: outputs, inputs, and internal states. Based on the design of our motor task, we assume that a movement on every trial goes through two phases: planning and execution. This is represented by two boxes seen in Figure 2a. The inputs of planning are speed and direction while the output is RT. The input of execution is perturbation as well as the RT from planning while the output is SE. Internal states are drawn as a black dashed line for illustrative purposes. They provide feedback for both planning and execution. This is because the internal states update is based on trial history (such as past performance or trial conditions). This information then flows through our system to affect the outputs. Though it is not labeled, the dotted line from planning to execution also carries the inputs from the planning system (speed and direction) to be available for the downstream system. Therefore speed and direction are also available for modeling SE. However, perturbation is not available for RT because perturbations happen after RT. However, the history of the trial conditions from previous trials is available through the internal states.

The outputs, RT and SE, are denoted 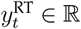 and 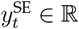 respectively. They are directly measured from the behavioral data during the motor task. The RT was normalized using the *z*-score before any modeling was performed so each subject followed a standard normal distribution (*i.e.*, 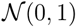). The SE innately followed a continuous uniform distribution (*i.e.*, 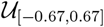) and was not normalized. Refer to Supplementary Table 5 for the statistics of outputs.

The inputs are the trial conditions of the motor task: speed, direction, and perturbation. They are also directly measured. They are described as categorical variables, denoting speed as 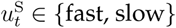, direction as 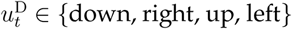, and perturbation as 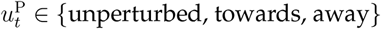.

The internal states we defined are the *error state* and *perturbed state*, denoted 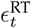 and 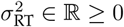 respectively.

Our system is broken down into the phases of planning and execution (Fig. 2a). The behavioral outputs of planning and execution are RT^[86]^ and SE^[25]^, respectively. They are separated by a delay for maximal separation^[87]^. It is important to note that the states remain constant between planning to execution since neither state has information to update until after a movement is complete. Each phase is associated with its own mathematical function relating the outputs as a linear combination of states and inputs available on trial *t*. The task begins with the planning phase, written as:

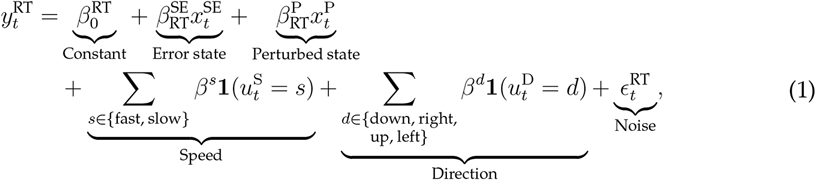

Where 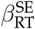 is an independent normal random variable with zero mean and variance 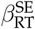. In other words, it defines the output of planning on trial *t* as RT and is the linear combination of a constant, error state, perturbed state, speed, and direction on trial *t*, scaled by their respective weights (*β*’s). This is followed by the execution phase written as:

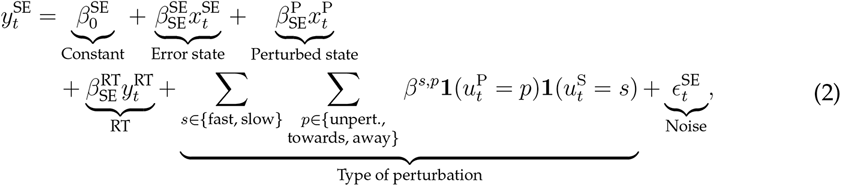

where 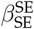 is an independent normal random variable with zero mean and variance 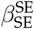. It defines the output of execution on trial *t* as SE and is the linear combination of a constant, error state, perturbed state, RT, and the combination of speed and perturbation on trial *t*, scaled by their respective weights (*β*’s). By our definition, RT will always be available as an input for SE. Though speed is not a direct input to execution (Fig. 2a), it also carries over from planning. We found the combination of speed and perturbation captured SE well as compared to any other linear combination of trial conditions. This combination is supported by intuition as well as in literature^[88]^. On one hand, a slow trial with an away perturbation could help subjects reduce the magnitude of their SE by forcing them to move slower. On the other hand, a fast trial with an away perturbation could make it harder to match the speed, making a negative SE (too slow) more believable.

To capture the history of their performance of speed error during the task, we used SE to update the error state:

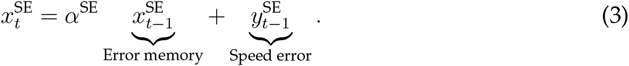

The degree to which the previous state weighs into the current state is scaled by *α*^SE^, which ranges between 0 and 1. An *α*^SE^ closer to 0 means the error state will quickly decay to 0 on subsequent trials while a *α*^SE^ closer to 1 means the error state will retain its value, such as the case for subjects who carry information over from trial-to-trial. The input is the SE from the previous trial. It ranges in value between −0.67 and 0.67, where a positive SE (*y*^SE^ *>* 0) means they moved slower than instructed and a negative SE (*y*^SE^ *<* 0) means they moved faster than instructed. Therefore, a positive error state indicates the accumulation of trials that were slower than instructed whereas a negative error state indicates the accumulation of trials that were faster than instructed. The sign of the weights *β*^SE^ in equations (1) and (2) depicts how the behavior of a subject would respond to the error state. Take the case when the error state is positive (*i.e.*, moving slower than instructed). A positive *β*^SE^ would increase the RT, thus subjects would react slower after trials in which they moved slower than instructed. Conversely, a negative *β*^SE^ would decrease their RT and subjects would react faster after trials in which they moved slower than instructed. For SE, a positive *β*^SE^ would increase the SE, meaning subjects would continue to move slower than instructed on subsequent trials. A negative *β*^SE^ would decrease the SE, meaning subjects would move faster than instructed on subsequent trials.

To capture the effect of perturbations on their behavior, we used an indicator on whether the perturbation input detected a perturbation either towards or away to update the perturbed state:

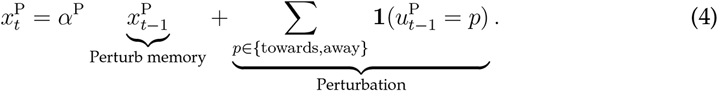

The perturbed state receives a positive pulse when perturbation was applied, regardless of the type of perturbation that was applied. Therefore, state only deviated from 0 when a perturbation was introduced. In the absence of a perturbation, the system would decay with a rate constant of *α*^P^, which ranges between 0 and 1 whereas a *α*^P^ closer to 0 indicates that the state will decay back to 0 by the next trial. A *α*^P^ closer to 1 means that subjects would carry over perturbations into subsequent trials through the perturbed state if it has not fully decayed to 0. Because of its structure, the perturbed state can only be positive or 0, where 0 means that perturbations do not effect behavior. The perturbed state effects behavior based on the sign of *β*^P^ in equations (1) and (2), so long as the perturbed state is not 0. In terms of RT, a positive 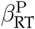 will increase the RT after a perturbation. Therefore, subjects with a positive weight will react slower than average after perturbation trials. A perturbation will also effect the SE based on the sign of 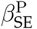. A positive 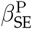 will SE increase SE after perturbation trials, *i.e.*, subjects will move slower than instructed after perturbations. Subjects where both weights are positive suggests that they hesitant in response to recent perturbations captured by their perturbed state.

Unlike the other elements in our system, the internal states cannot be directly measured because they are subjective. They are an internal representation of the environment that an individual defines which evolves given new information. Instead, internal states are dynamically updated on a trial-by-trial basis by weighing their past states. Both must be estimated using a first-order state evolution equation, whose function is controlled by what is added to it in addition to their past states. The general solution is:

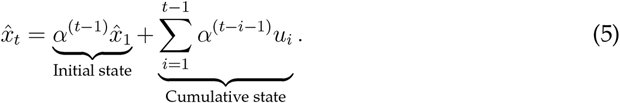

Therefore, the states are not simply a weight of the previous trial but capture the accumulation of history from previous trials.

Thus, Equations (3) to (2) make up our system. But the system is not complete until fitting the models. Model fitting consisted of finding the combination of weights (*α*’s and *β*’s) that minimized the root-mean-squared error between the observed and estimated outputs for each subject using all complete trials. First, the *α*’s were found using a grid search between 0.01 and 0.99 at an interval of 0.01 with the initial conditions 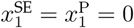 to estimate the internal states. Additionally, the internal states were normalized using the *z*-score so their weights could be compared across subjects. Then, the weights were found using methods about generalized linear model, which solves for the maximum likelihood estimation. The resulting states and weights were applied to estimate the outputs. The combination of weights with the largest Pearson correlation value between observed and estimated outputs was selected as the final model. This process was repeated for each subject to create their custom model fitting, complete with their own individual evolution of internal states.

To show the adequacy of our model, we also built a subject-specific linear model that relied only on trial conditions. The linear model was a simple linear regression between the outputs (*i.e.*, RT and SE) and inputs (*i.e.*, speed, direction and RT, speed, perturbation).All models were evaluated using the coefficient of determination and deviance for comparison. The coefficient of determination was used to quantify how much the variability of the outputs can be explained by the inputs. It ranges in value between 0 and 1, where 1 means that the estimated outputs completely accounts for all the variability of the observed outputs. The deviance was used to compare the error of each model. This can be helpful as an absolute measurement of goodness-of-fit to compare models by measuring the trade-off between model complexity and goodness-of-fit. Deviance can be any positive value, where a deviance of 0 means that the model describes the observed outputs perfectly.

### Neural data preprocessing

We used spectral analysis to preprocess the LFP activity from each channel. First, a Notch filter was applied using notches located at the fundamental frequency of 60 Hz with bandwidth at the −1 dB point set to 3 Hz. Next, the oscillatory power was calculated using a continuous wavelet transform with a logarithmic scale vector ranging 1–200 Hz and complex Morlet wavelet with a default radian frequency of *ω*_0_ = 6. The resulting instantaneous power spectral density was divided into overlapping time bins using a window of 100 ms every 50 ms. All overlapping time bins were averaged together and labeled with the last time index corresponding to that window. Finally, the averaged power spectral density was normalized to equally weigh all frequency bins by taking the *z*-score of the natural logarithm of the power in each frequency bin over the entire recording session time. All channel recordings were visually inspected for artifacts before and after preprocessing. Examples of artifacts include broadband effects, abnormal bursts of power, and faulty recordings. Any channels with artifacts were disregarded for the entire session. Figure 4c shows the result of the spectral analysis for a channel as a spectrogram, where the color of each pixel represents the normalized power indexed by the color bar (between −3 and 3) at a specified frequency and time.

The results of our analysis depend heavily on how channels are aggregated using their anatomical labels. Therefore, it was important to ensure that their labels were unbiased across subjects. We applied a semi-automated electrode localization protocol to determine the coordinates of each channels per subject by fusing their preoperative MRI with postoperative CT^[89]^. This protocol also labeled each channel using an anatomical atlas^[90]^ and a large-scale functional brain network atlas^[91]^ based on their coordinates onto a subject-specific cortical parcellation^[92]^. The anatomical labels were validated by a clinician. To visualize the coverage of channels across the populations, their coordinates were warped from the subject’s native space to the standard Montreal Neurological Institute (MNI) atlas space^[93]^. All channels could then be visualized on a common template brain (`cvs avg35 inMNI152`) from FreeSurfer^[92]^. Figure 4a shows the electrode placement of all subjects projected onto this template brain.

### Identifying regions that encode internal states

After neural data preprocessing, we used an unsupervised paradigm to identify where internal states are encoded in the brain of the population. We accomplished this using a non-parametric cluster statistic^[35]^. Details on how this method was applied to similar data by our group^[19]^. Here, we used a two-tailed permutation test with *N* = 1, 000 and a significance threshold of *α* = 0.05. In short, the procedure works by finding windows of time and frequency where the LFP in a channel (measured as power in the spectrogram) that covary with an internal state across the population, which are formed by aggregating the anatomical labels of the channels from all subjects. The result are windows of time and frequency, known as *clusters*, across brain regions.

Clusters that were too small (*i.e.*, had windows less than 250 ms in time, one octave in frequency, or area smaller than the minimum time and frequency windows specified) were discarded. We also discarded regions that had less than two subjects contributing to the cluster. A false discovery rate of *q* = 0.015 was then applied to correct for multiple comparisons between regions and epochs. Clusters are confined to predefined windows of time set by the epoch it was recorded from for the analysis. However, since neural activity is continuous, information it may be encoding could carry over from one epoch to the next. Therefore, clusters in the same region with overlapping frequency bins across epochs were grouped for further analysis. This also meant that a region could come up multiple times, such as the case if separate clusters were found in different frequency bins in the same epoch.

Each group of clusters was assigned a frequency band. The *frequency band* was identified by matching the frequency bins to frequency bands commonly defined in literature: delta (1–4 Hz), theta (4–8 Hz), alpha (8–15 Hz), beta (15–30 Hz), low gamma (30–60 Hz), high gamma (60–100 Hz), and hyper gamma (100–200 Hz). If the group of clusters spanned multiple frequency bands, then the band that made up the majority of the group of clusters was prioritized.

Further, we observed two distinct temporal patterns of activity that described each group of clusters; persistent or phasic. *Persistent activity* refers to a group of clusters whose activity stretched across all epochs during a trial. *Phasic activity* refers to a group of clusters whose activity only appeared during specific epochs of a trial related the movement phase (*i.e.*, planning, execution, feedback).

### Calculating encoding strength

We hypothesized that neural activity in encoding networks of top performers will modulate with internal states more than bottom performers. To test this, we quantified how well a region covaries with the internal state using encoding strength and compared it to session performance across subjects. We calculated the encoding strength of group of clusters for each subject by replicating the procedure from the non-parametric cluster statistic. For each group of clusters, we first averaged the neural activity in its time-frequency window across the epochs it spans for each channel, trial, and subject. Next, we then found the magnitude of the Spearman correlation value between this averaged neural activity and the internal state across all trials for each channel and subject. Finally, we averaged these correlation magnitudes across channels in a subject with the same group of clusters. This value is the *encoding strength* of the group of clusters for a subject of a region. Because it is derived from the magnitude of the correlation, it takes on a value between 0 and 1, where 1 means the neural activity exactly follows the internal state across trials.

To identify regions that encode internal states and performance, we correlated the encoding strength of each group of clusters to the session performance across subjects. This allowed us to identify the regions that not only encoded the internal states, but also related to session performance (*i.e.*, subject variability). These performance-related regions were those whose Pearson correlation value exceeded 0.75 or *p*-value was significant (*p <* 0.05). We did not rely solely on significant *p*-values because the largest possible sample size (*i.e.*, *n ≤* 10) was small. Further, we used the magnitude of correlation value because we were interested in the relationship between encoding strength and session performance, not the direction of encoding (*i.e.*, positive or negative). Because we were focused on comparing top and bottom performers, we rejected regions that did not have both type of performers.

A table of these regions can be found in Table 2 and Table 3 for error and perturbed state, respectively. They are also displayed on an inflated brain template in Figure 5a and Figure 5b, where gyri and sulci can be visualized together.

### Calculating connectivity strength

We hypothesized that differences in functional connectivity within and between networks that encode internal states account for learning strategies of top and bottom performers. To test this, we compared functional connectivity with session performance across subjects. Functional connectivity is defined as dynamic connections between neuronal populations through oscillatory activity^[94]^. There are many ways to calculate functional connectivity^[94]^. We chose to use cross-correlation using a lag of 0, which simply becomes the Pearson correlation value^[36, 95]^.

To calculate connectivity, we began by averaged the neural activity of each group of clusters using its time-frequency window across the epochs it spans for each channel, trial, and subject. Then, we found the magnitude of the Pearson correlation value between each pair the averaged neural activity of group of clusters and channels for each subject across trials. Pairs of channels in the same group of clusters were then averaged within each subject. These values represent the *connectivity strength* of the pair of regions for each subject. Because it is derived from the magnitude of the correlation, it takes on a value between 0 and 1, where 1 means the activity of the pair is perfectly correlated across trials.

To combine the correlation values across subjects for a population analysis, correlation value were first transformed using Fischer’s *z* transformation^[96]^ before averaging between the same pair of region labels. After averaging, the values were transformed back into correlation values using Fischer’s *z* transformation^[96]^ due to our small sample size^[97]^. These values represent the connectivity strength of the population. Any pairs that had fewer than two subjects were excluded. Likewise, connectivity strength on the diagonal (*i.e.*, autocorrelations) were ignored for both subject and population connectivity strengths.

To identify pairs of regions whose connectivity encodes the internal states and performance, we correlated the connectivity strength of each pair of regions to the session performance across subjects. Following the same criteria as encoding strength, only pairs whose Pearson correlation value exceeded 0.75 or *p*-value was significant (*p <* 0.05) were considered. Any pairs that did not include both a top and bottom performer were excluded.

## Supplementary Information

### Supplementary Figures

**Supplementary Figure 1.**
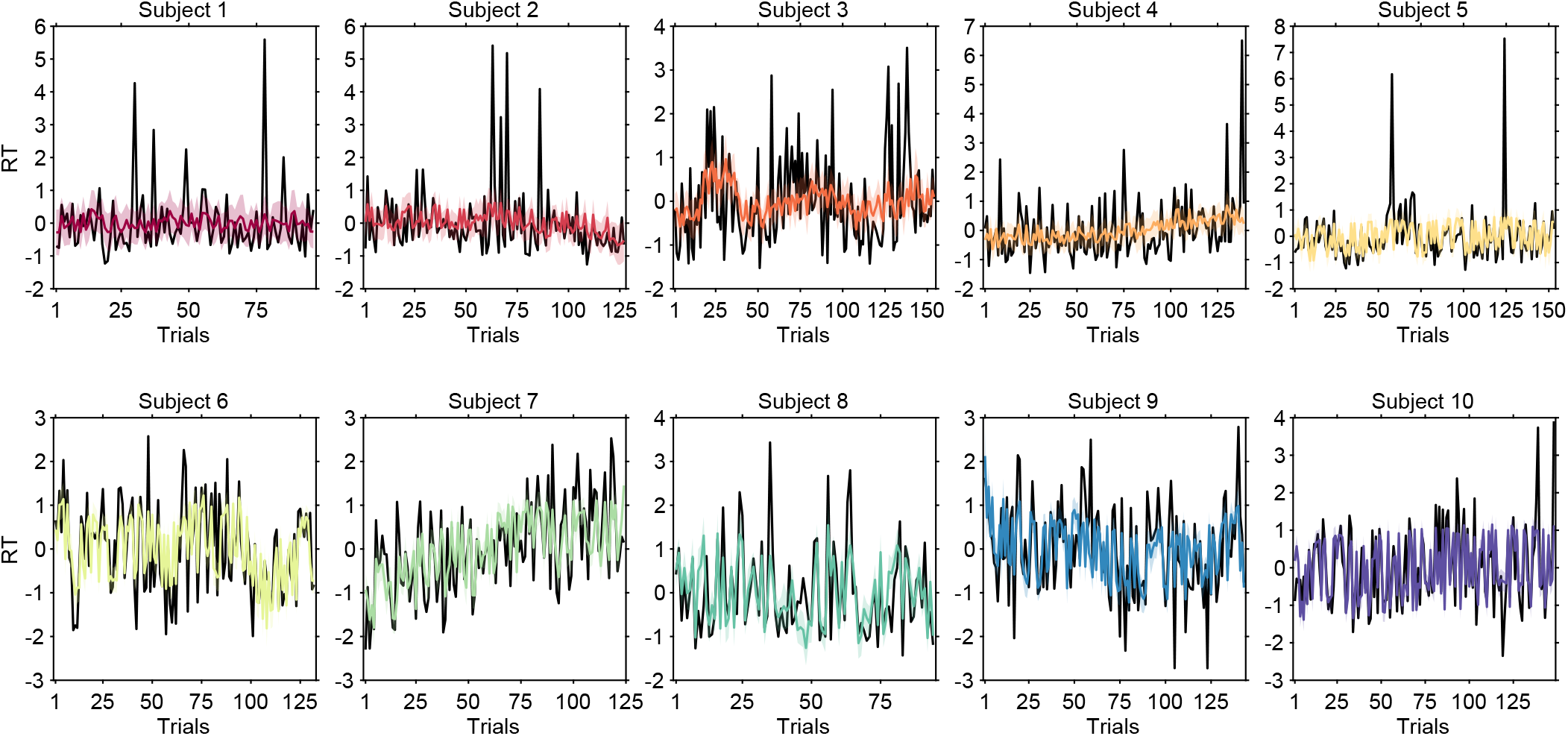
Estimation of RT. Overlay of observed RT (black solid line) and estimated RT (solid color line) across all trials. The RT of each subject was normalized using the *z*-score before fitting and plotting the models. Each subject is color-coded in their panel. The black solid line represents the observed RT. The solid color line represents the estimated RT and the shaded color represents the 95 % confidence bound.

**Supplementary Figure 2.**
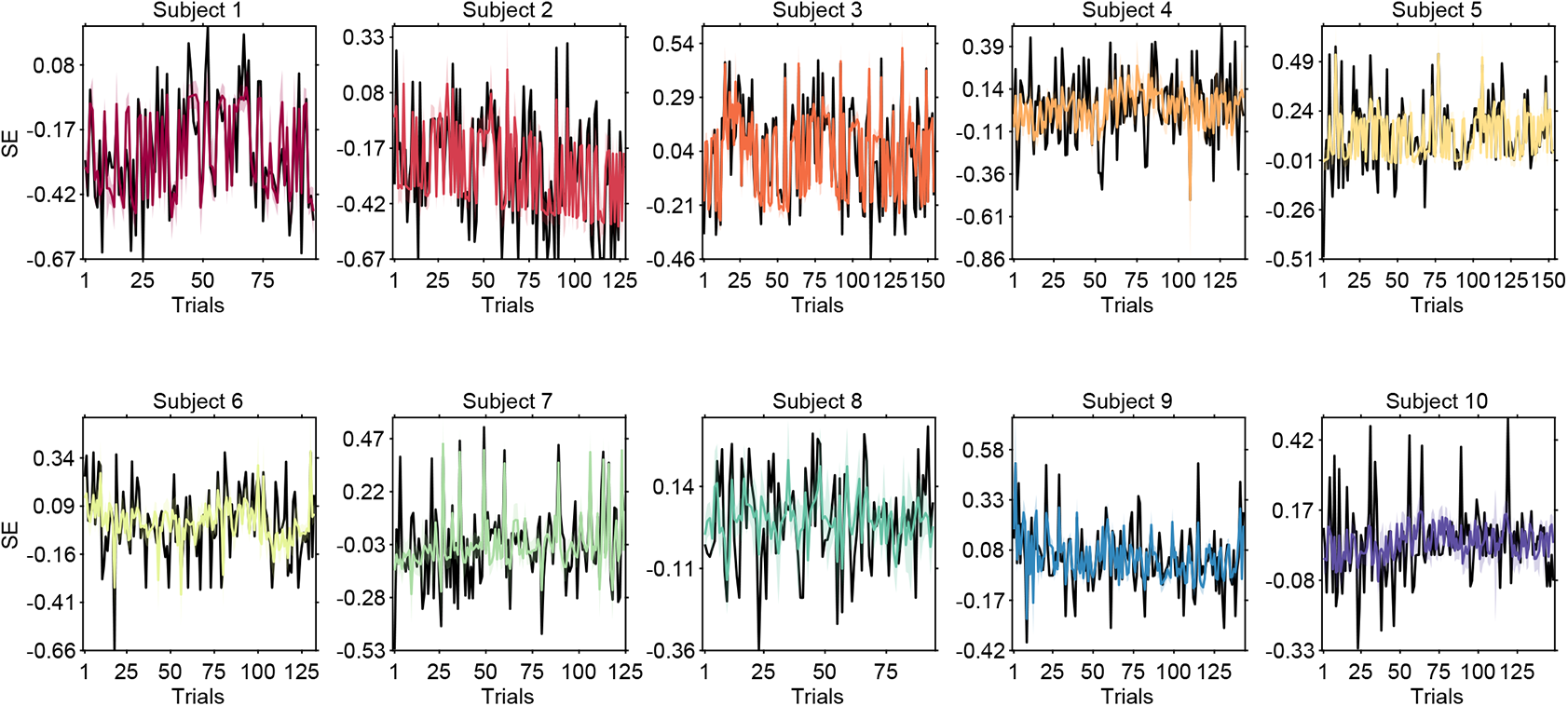
Estimation of SE. Overlay of observed SE (black solid line) and estimated SE (solid color line) across all trials. Each subject is color-coded in their panel. The black solid line represents the observed SE. The solid color line represents the estimated SE and the shaded color represents the 95 % confidence bound.

**Supplementary Figure 3.**
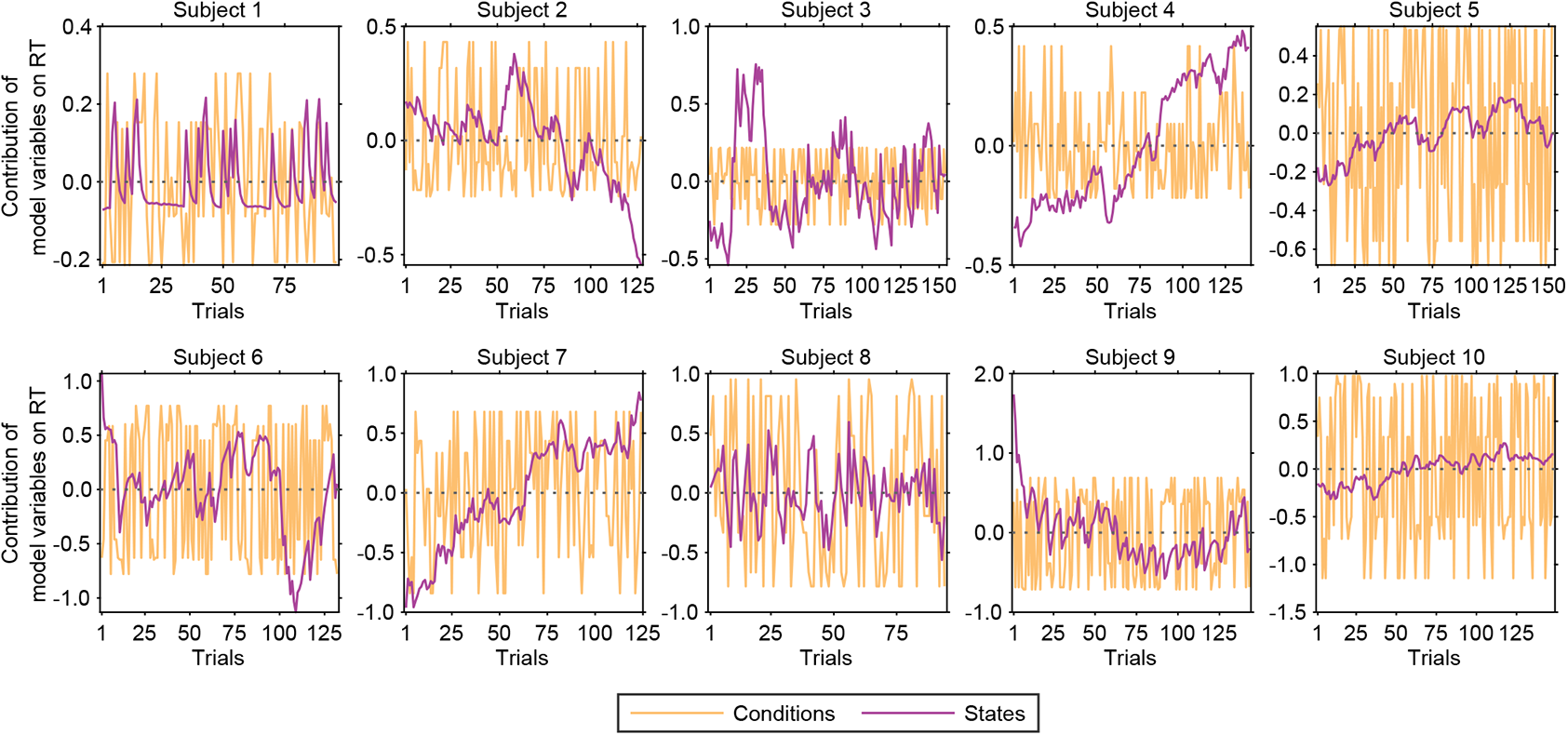
Contribution of model variables to RT. Contribution of the internal states (purple solid line) and trial conditions (orange solid line) in the RT model across trials.

**Supplementary Figure 4.**
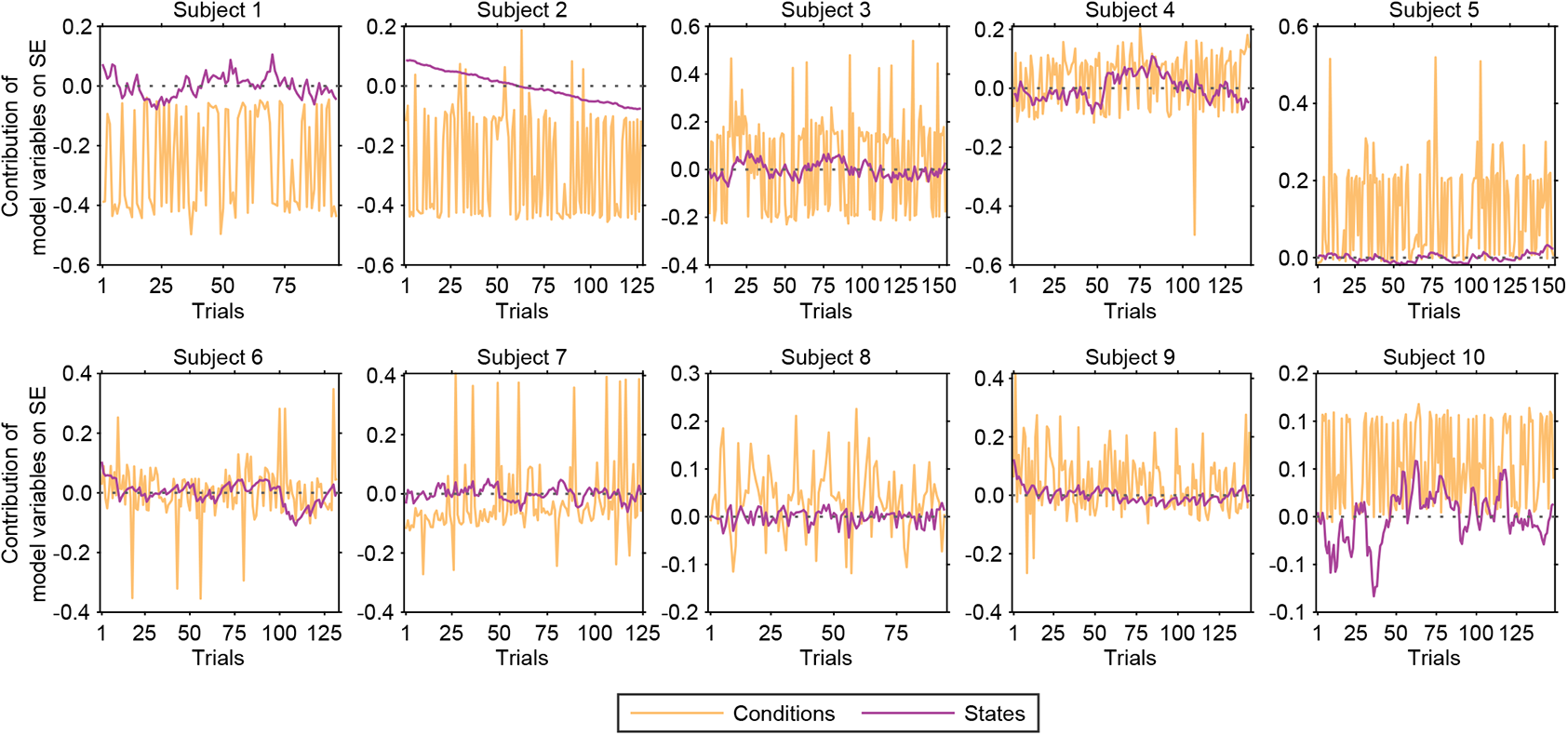
Contribution of model variables to SE. Contribution of the internal states (purple solid line) and trial conditions (orange solid line) in the SE model across trials.

**Supplementary Figure 5.**
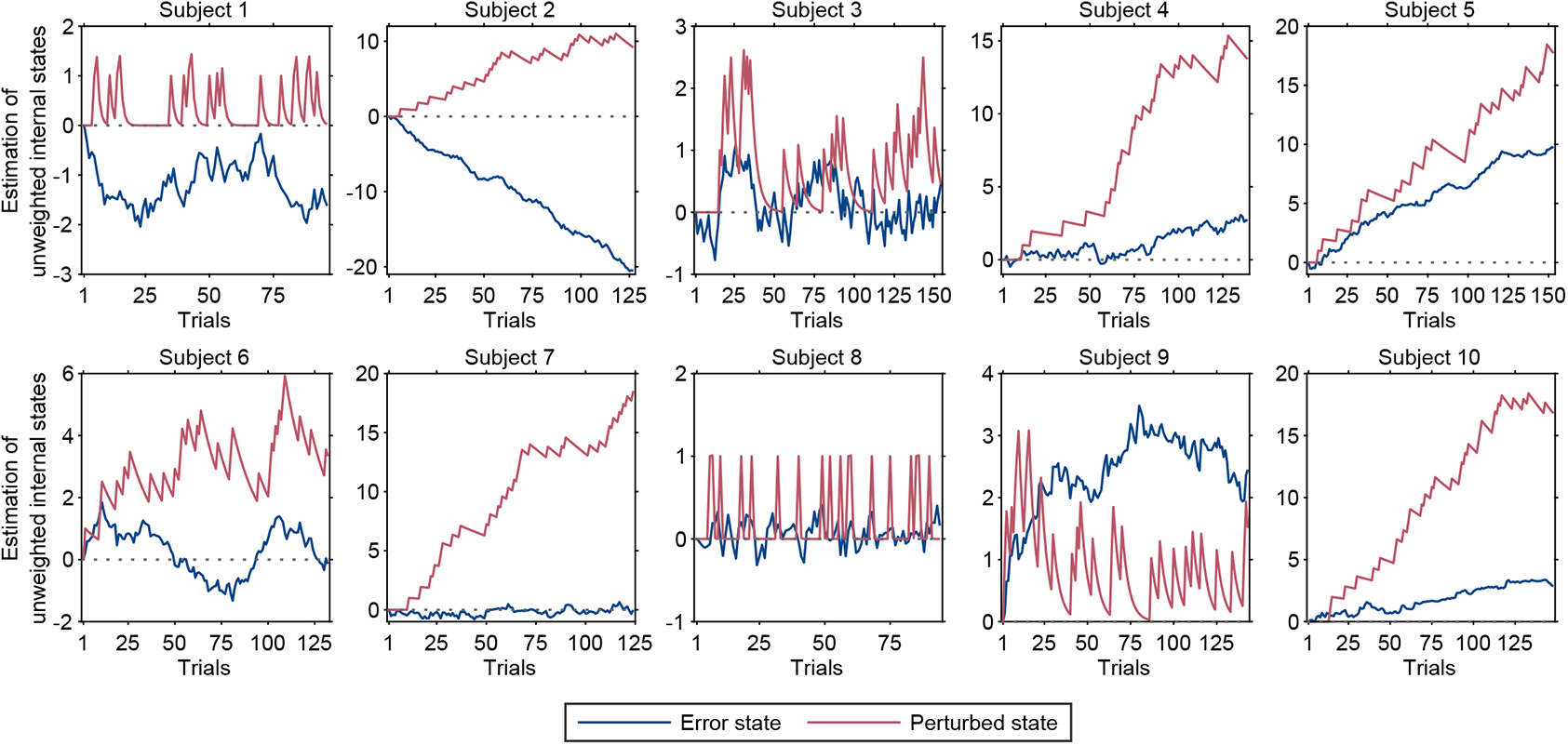
Estimation of internal states. Estimation of error state (blue solid line) and perturbed state (red solid line) over trials for each subject. The states were used to estimate both RT and SE by fitting each model with different weights. The y-axis here does not reflect the actual values used in equations (1) and (2) because the internal states were normalized using the *z*-score before fitting.

**Supplementary Figure 6.**
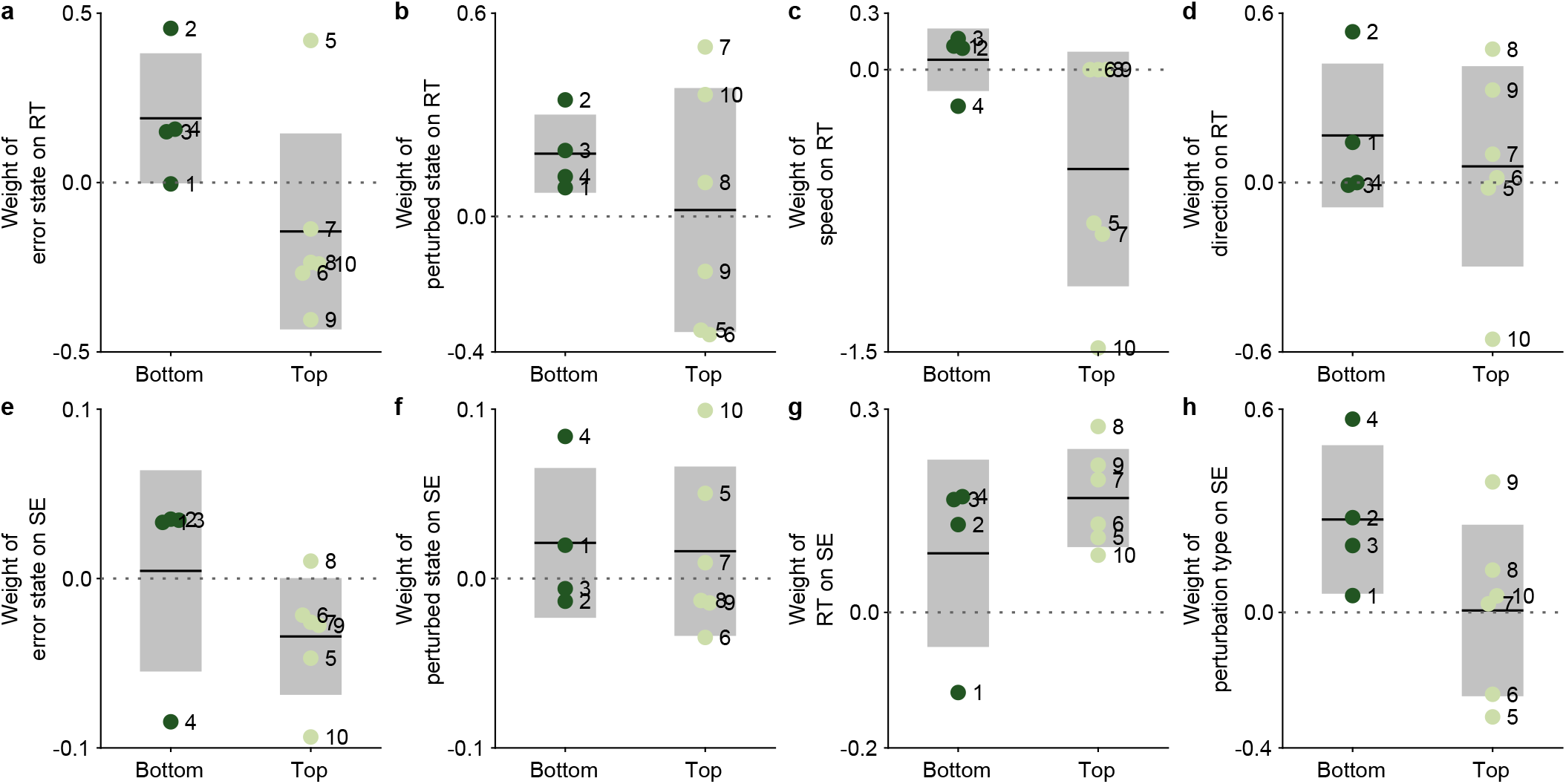
Comparison between model weights and session performance. Markers are color-coded for each subject. Weight of **a,** error state, **b,** perturbed state, **c,** speed, **d,** and direction on RT in equation (1). Weight of **e,** error state, **f,** perturbed state, **g,** RT, **h,** and perturbation type on SE in equation (2).

### Supplementary Tables

**Supplementary Table 1.** Number of completed trials for each trial conditions (*e.g.*, speed, direction, perturbation) and session performance (as a percent) for each subject divided into top and bottom performers using the average population session performance (51 %) as a threshold.

| Subject | Completed | Number of trials |  |  |  |  |  |  |  |  | Session performance (%) | Performer group |
| --- | --- | --- | --- | --- | --- | --- | --- | --- | --- | --- | --- | --- |
|  |  | Speed<br>fast slow |  | Direction<br>down right up left |  |  |  | Perturbation<br>unpert. towards away |  |  |  |  |
| 1 | 96 | 45 | 51 | 27 | 20 | 26 | 23 | 76 | 8 | 12 | 24 | Bottom |
| 2 | 127 | 53 | 74 | 43 | 39 | 34 | 30 | 102 | 10 | 15 | 29 |  |
| 3 | 154 | 81 | 73 | 37 | 38 | 43 | 36 | 128 | 6 | 20 | 30 |  |
| 4 | 139 | 77 | 62 | 39 | 36 | 29 | 35 | 115 | 8 | 16 | 50 |  |
| 5 | 153 | 68 | 85 | 36 | 45 | 39 | 33 | 122 | 7 | 24 | 53 | Top |
| 6 | 132 | 64 | 68 | 37 | 30 | 33 | 32 | 105 | 13 | 14 | 56 |  |
| 7 | 124 | 56 | 68 | 38 | 30 | 34 | 22 | 94 | 8 | 22 | 59 |  |
| 8 | 94 | 53 | 41 | 16 | 29 | 20 | 29 | 76 | 6 | 12 | 63 |  |
| 9 | 143 | 65 | 78 | 46 | 29 | 36 | 32 | 113 | 3 | 27 | 66 |  |
| 10 | 148 | 74 | 74 | 31 | 43 | 39 | 35 | 117 | 7 | 24 | 81 |  |

**Supplementary Table 2.** Statistics for the reaction time and speed error for each subject using all of their completed trials.

| Subject | Reaction time (s) |  | Speed error |  |
| --- | --- | --- | --- | --- |
|  | Mean | STD | Mean | STD |
| 1 | 0.98 | 0.26 | -0.26 | 0.20 |
| 2 | 0.80 | 0.18 | -0.26 | 0.24 |
| 3 | 0.91 | 0.19 | 0.03 | 0.22 |
| 4 | 0.93 | 0.22 | 0.03 | 0.20 |
| 5 | 0.66 | 0.20 | 0.12 | 0.17 |
| 6 | 1.07 | 0.27 | 0.0 | 0.18 |
| 7 | 0.73 | 0.12 | -0.03 | 0.19 |
| 8 | 0.80 | 0.27 | 0.04 | 0.15 |
| 9 | 0.67 | 0.18 | 0.04 | 0.15 |
| 10 | 0.52 | 0.11 | 0.05 | 0.13 |

**Supplementary Table 3.**
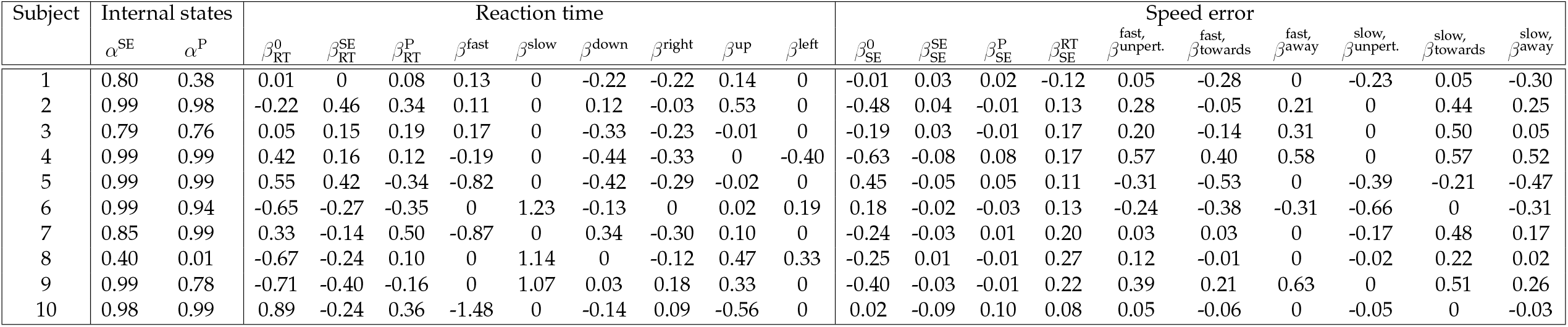
Weights of model for each subject used in equations (3) and (4) for *α*’s and equations (1) and (2) for *β*’s.

## References

1. Faisal, A. A., Selen, L. P. J. & Wolpert, D. M. Noise in the nervous system. Nature Reviews Neuroscience 9, 292–303 (2008).

2. Wolpert, D. M., Diedrichsen, J. & Flanagan, J. R. Principles of sensorimotor learning. Nature Reviews Neuroscience 12, 739–751 (2011).

3. Dhawale, A. K., Smith, M. A. & Ölveczky, B. P. The role of variability in motor learning. Annual Review of Neuroscience 40, 479–498 (2017).

4. Ölveczky, B. P., Andalman, A. S. & Fee, M. S. Vocal experimentation in the juvenile songbird requires a basal ganglia circuit. PLoS Biology 3, e153 (2005).

5. Wu, H. G., Miyamoto, Y. R., Castro, L. N. G., Ölveczky, B. P. & Smith, M. A. Temporal structure of motor variability is dynamically regulated and predicts motor learning ability. Nature Neuroscience 17, 312–321 (2014).

6. Mir, P. et al. Motivation and movement: the effect of monetary incentive on performance speed. Experimental Brain Research 209, 551–559 (2011).

7. Manohar, S. G., Muhammed, K., Fallon, S. J. & Husain, M. Motivation dynamically increases noise resistance by internal feedback during movement. Neuropsychologia (2018).

8. Galaro, J. K., Celnik, P. & Chib, V. S. Motor cortex excitability reflects the subjective value of reward and mediates its effects on incentive-motivated performance. Journal of Neuroscience 39, 1236–1248 (2019).

9. Kiani, R. & Shadlen, M. N. Representation of confidence associated with a decision by neurons in the parietal cortex. science 324, 759–764 (2009).

10. Zylberberg, A., Fetsch, C. R. & Shadlen, M. N. The influence of evidence volatility on choice, reaction time and confidence in a perceptual decision. eLife 5, e17688 (2016).

11. Schmidt, L. et al. Get aroused and be stronger: Emotional facilitation of physical effort in the human brain. Journal of Neuroscience 29, 9450–9457 (2009).

12. Lawrence, G. P., Khan, M. A. & Hardy, L. The effect of state anxiety on the online and offline control of fast target-directed movements. Psychological Research 77, 422–433 (2012).

13. Blakemore, R. L. & Vuilleumier, P. An emotional call to action: Integrating affective neuroscience in models of motor control. Emotion Review 9, 299–309 (2016).

14. van Den Berg, R. et al. A common mechanism underlies changes of mind about decisions and confidence. eLife 5, e12192 (2016).

15. Braun, A., Urai, A. E. & Donner, T. H. Adaptive history biases result from confidence-weighted accumulation of past choices. Journal of Neuroscience 38, 2418–2429 (2018).

16. Calhoun, A. J., Pillow, J. W. & Murthy, M. Unsupervised identification of the internal states that shape natural behavior. Nature Neuroscience 22, 2040–2049 (2019).

17. Ashwood, Z. C. et al. Mice alternate between discrete strategies during perceptual decision-making. Nature Neuroscience 25, 201–212 (2022).

18. Gründemann, J. et al. Amygdala ensembles encode behavioral states. Science 364, eaav8736 (2019).

19. Sacré, P. et al. Risk-taking bias in human decision-making is encoded via a right–left brain push–pull system. Proceedings of the National Academy of Sciences 116, 1404–1413 (2019).

20. Hwang, E. J., Dahlen, J. E., Mukundan, M. & Komiyama, T. History-based action selection bias in posterior parietal cortex. Nature Communications 8, 1–14 (2017).

21. Wei, Z., Inagaki, H., Li, N., Svoboda, K. & Druckmann, S. An orderly single-trial organization of population dynamics in premotor cortex predicts behavioral variability. Nature Communications 10, 1–14 (2019).

22. Cowley, B. R. et al. Slow drift of neural activity as a signature of impulsivity in macaque visual and prefrontal cortex. Neuron 108, 551–567 (2020).

23. Wolpert, D. M. & Landy, M. S. Motor control is decision-making. Current opinion in neurobiology 22, 996–1003 (2012).

24. Churchland, M. M., Afshar, A. & Shenoy, K. V. A central source of movement variability. Neuron 52, 1085–1096 (2006).

25. van Beers, R. J., Haggard, P. & Wolpert, D. M. The role of execution noise in movement variability. Journal of Neurophysiology 91, 1050–1063 (2004).

26. Renart, A. & Machens, C. K. Variability in neural activity and behavior. Current Opinion in Neurobiology 25, 211–220 (2014).

27. Tourangeau, R. & Rasinski, K. A. Cognitive processes underlying context effects in attitude measurement. Psychological Bulletin 103, 299 (1988).

28. Critchley, H. D., Elliott, R., Mathias, C. J. & Dolan, R. J. Neural activity relating to generation and representation of galvanic skin conductance responses: a functional magnetic resonance imaging study. Journal of Neuroscience 20, 3033–3040 (2000).

29. Lane, R. D. et al. Neural correlates of heart rate variability during emotion. NeuroImage 44, 213–222 (2009).

30. Thayer, J. F., Åhs, F., Fredrikson, M., Sollers III, J. J. & Wager, T. D. A meta-analysis of heart rate variability and neuroimaging studies: implications for heart rate variability as a marker of stress and health. Neuroscience & Biobehavioral Reviews 36, 747–756 (2012).

31. Podvalny, E., King, L. E. & He, B. J. Spectral signature and behavioral consequence of spontaneous shifts of pupil-linked arousal in human. eLife 10 (2021).

32. Schwarz, N. & Oyserman, D. Asking questions about behavior: Cognition, communication, and questionnaire construction. The American Journal of Evaluation 22, 127–160 (2001).

33. Kanwal, J. K. et al. Internal state: dynamic, interconnected communication loops distributed across body, brain, and time. Integrative and Comparative Biology (2021).

34. Logothetis, N. K. What we can do and what we cannot do with fMRI. Nature 453, 869–878 (2008).

35. Maris, E. & Oostenveld, R. Nonparametric statistical testing of EEG- and MEG-data. Journal of Neuroscience Methods 164, 177–190 (2007).

36. Friston, K. J. Functional and effective connectivity in neuroimaging: a synthesis. Human Brain Mapping 2, 56–78 (1994).

37. Harris, C. M. & Wolpert, D. M. Signal-dependent noise determines motor planning. Nature 394, 780–784 (1998).

38. Todorov, E. & Jordan, M. I. Optimal feedback control as a theory of motor coordination. Nature Neuroscience 5, 1226–1235 (2002).

39. Fine, M. S. & Thoroughman, K. A. Trial-by-trial transformation of error into sensorimotor adaptation changes with environmental dynamics. Journal of Neurophysiology 98, 1392–1404 (2007).

40. Fine, M. S. & Thoroughman, K. A. Motor adaptation to single force pulses: Sensitive to direction but insensitive to within-movement pulse placement and magnitude. Journal of Neurophysiology 96, 710–720 (2006).

41. Heitz, R. P. The speed-accuracy tradeoff: history, physiology, methodology, and behavior. Frontiers in Neuroscience 8, 150 (2014).

42. Steinhauser, M. & Yeung, N. Error awareness as evidence accumulation: effects of speed-accuracy trade-off on error signaling. Frontiers in Human Neuroscience 6, 240 (2012).

43. Van Veen, V., Krug, M. K. & Carter, C. S. The neural and computational basis of controlled speed-accuracy tradeoff during task performance. Journal of Cognitive Neuroscience 20, 1952–1965 (2008).

44. Agam, Y. et al. Network dynamics underlying speed-accuracy trade-offs in response to errors. PLoS One 8, e73692 (2013).

45. Eryurek, K. et al. Default mode and dorsal attention network involvement in visually guided motor sequence learning. Cortex 146, 89–105 (2022).

46. Fuster, J. M. & Alexander, G. E. Neuron activity related to short-term memory. Science 173, 652–654 (1971).

47. Lachaux, J.-P., Axmacher, N., Mormann, F., Halgren, E. & Crone, N. E. High-frequency neural activity and human cognition: past, present and possible future of intracranial eeg research. Progress in Neurobiology 98, 279–301 (2012).

48. Constantinidis, C. et al. Persistent spiking activity underlies working memory. Journal of Neuroscience 38, 7020–7028 (2018).

49. Gnadt, J. W. & Andersen, R. A. Memory related motor planning activity in posterior parietal cortex of macaque. Experimental Brain Research 70, 216–220 (1988).

50. Fiehler, K. et al. Working memory maintenance of grasp-target information in the human posterior parietal cortex. NeuroImage 54, 2401–2411 (2011).

51. Noudoost, B., Clark, K. L. & Moore, T. Working memory gates visual input to primate prefrontal neurons. eLife 10, e64814 (2021).

52. Majerus, S., Péters, F., Bouffier, M., Cowan, N. & Phillips, C. The dorsal attention network reflects both encoding load and top–down control during working memory. Journal of Cognitive Neuroscience 30, 144–159 (2018).

53. Bledowski, C., Rahm, B. & Rowe, J. B. What “works” in working memory? separate systems for selection and updating of critical information. Journal of Neuroscience 29, 13735–13741 (2009).

54. Seidler, R. D., Bo, J. & Anguera, J. A. Neurocognitive contributions to motor skill learning: the role of working memory. Journal of Motor Behavior 44, 445–453 (2012).

55. Masse, N. Y., Rosen, M. C. & Freedman, D. J. Reevaluating the role of persistent neural activity in short-term memory. Trends in Cognitive Sciences 24, 242–258 (2020).

56. Robertson, I. H., Manly, T., Andrade, J., Baddeley, B. T. & Yiend, J. ’oops!’: performance correlates of everyday attentional failures in traumatic brain injured and normal subjects. Neuropsychologia 35, 747–758 (1997).

57. Pessoa, L., Gutierrez, E., Bandettini, P. A. & Ungerleider, L. G. Neural correlates of visual working memory: fmri amplitude predicts task performance. Neuron 35, 975–987 (2002).

58. Padilla, M. L., Wood, R. A., Hale, L. A. & Knight, R. T. Lapses in a prefrontal-extrastriate preparatory attention network predict mistakes. Journal of Cognitive Neuroscience 18, 1477–1487 (2006).

59. Fortenbaugh, F. C., Rothlein, D., McGlinchey, R., DeGutis, J. & Esterman, M. Tracking behavioral and neural fluctuations during sustained attention: A robust replication and extension. NeuroImage 171, 148–164 (2018).

60. Kucyi, A. et al. Electrophysiological dynamics of antagonistic brain networks reflect attentional fluctuations. Nature Communications 11, 1–14 (2020).

61. Esterman, M., Noonan, S. K., Rosenberg, M. & DeGutis, J. In the zone or zoning out? tracking behavioral and neural fluctuations during sustained attention. Cerebral Cortex 23, 2712–2723 (2013).

62. McVay, J. C. & Kane, M. J. Drifting from slow to “d’oh!”: Working memory capacity and mind wandering predict extreme reaction times and executive control errors. Journal of Experimental Psychology: Learning, Memory, and Cognition 38, 525 (2012).

63. Adam, K. C., Mance, I., Fukuda, K. & Vogel, E. K. The contribution of attentional lapses to individual differences in visual working memory capacity. Journal of Cognitive Neuroscience 27, 1601–1616 (2015).

64. DeBettencourt, M. T., Keene, P. A., Awh, E. & Vogel, E. K. Real-time triggering reveals concurrent lapses of attention and working memory. Nature Human Behaviour 3, 808–816 (2019).

65. Machner, B. et al. Resting-state functional connectivity in the dorsal attention network relates to behavioral performance in spatial attention tasks and may show task-related adaptation. Frontiers in human neuroscience 15, 757128 (2022).

66. Roberts, S. D. et al. Investigation of baseline attention, executive control, and performance variability in female varsity athletes. Brain Imaging and Behavior 1–10 (2022).

67. Burgess, G. C. et al. Attentional control activation relates to working memory in attention-deficit/hyperactivity disorder. Biological psychiatry 67, 632–640 (2010).

68. Salmi, J. et al. Out of focus–Brain attention control deficits in adult ADHD. Brain Research 1692, 12–22 (2018).

69. Brandman, T., Malach, R. & Simony, E. The surprising role of the default mode network in naturalistic perception. Communications Biology 4, 1–9 (2021).

70. Kerr, M. S. D. et al. The role of associative cortices and hippocampus during movement perturbations. Frontiers in Neural Circuits 11, 26 (2017).

71. Dohmatob, E., Dumas, G. & Bzdok, D. Dark control: The default mode network as a reinforcement learning agent. Human Brain Mapping 41, 3318–3341 (2020).

72. Zhang, H. et al. Motor imagery learning modulates functional connectivity of multiple brain systems in resting state. PLoS One 9, e85489 (2014).

73. Albouy, G. et al. Neural correlates of performance variability during motor sequence acquisition. NeuroImage 60, 324–331 (2012).

74. Sali, A. W., Courtney, S. M. & Yantis, S. Spontaneous fluctuations in the flexible control of covert attention. Journal of Neuroscience 36, 445–454 (2016).

75. Hinds, O. et al. Roles of default-mode network and supplementary motor area in human vigilance performance: evidence from real-time fmri. Journal of Neurophysiology 109, 1250–1258 (2013).

76. Hsu, H. M., Yao, Z.-F., Hwang, K. & Hsieh, S. Between-module functional connectivity of the salient ventral attention network and dorsal attention network is associated with motor inhibition. PLoS One 15, e0242985 (2020).

77. Diedrichsen, J., Hashambhoy, Y., Rane, T. & Shadmehr, R. Neural correlates of reach errors. Journal of Neuroscience 25, 9919–9931 (2005).

78. Cléry-Melin, M.-L. et al. Why don’t you try harder? An investigation of effort production in major depression. PloS one 6, e23178 (2011).

79. Burris, K. et al. Sensorimotor abilities predict on-field performance in professional baseball. Scientific Reports 8, 1–9 (2018).

80. González-Martínez, J. et al. Technique, results, and complications related to robot-assisted stereoelectroencephalography. Neurosurgery 78, 169–180 (2015).

81. Johnson, M. A. et al. Performing behavioral tasks in subjects with intracranial electrodes. Journal of Visualized Experiments e51947 (2014).

82. Breault, M. S., Sacré, P., González-Martínez, J., Gale, J. T. & Sarma, S. V. An exploratory data analysis method for identifying brain regions and frequencies of interest from large-scale neural recordings. Journal of Computational Neuroscience (2018).

83. Breault, M. S. et al. Non-motor brain regions in non-dominant hemisphere dominate in decoding movement speed. Frontiers in Neuroscience 13, 715 (2019).

84. Asaad, W. F., Santhanam, N., McClellan, S. & Freedman, D. J. High performance execution of psychophysical tasks with complex visual stimuli in MATLAB. Journal of Neurophysiology 109, 249–260 (2013).

85. Asaad, W. F. & Eskandar, E. N. A flexible software tool for temporally-precise behavioral control in Matlab. Journal of Neuroscience Methods 174, 245–258 (2008).

86. Afshar, A. et al. Single-trial neural correlates of arm movement preparation. Neuron 71, 555–564 (2011).

87. Crammond, D. J. & Kalaska, J. F. Prior information in motor and premotor cortex: Activity during the delay period and effect on pre-movement activity. Journal of Neurophysiology 84, 986–1005 (2000).

88. Cluff, T. & Scott, S. H. Apparent and actual trajectory control depend on the behavioral context in upper limb motor tasks. Journal of Neuroscience 35, 12465–12476 (2015).

89. Stolk, A. et al. Integrated analysis of anatomical and electrophysiological human intracranial data. Nature Protocols 13, 1699–1723 (2018).

90. Destrieux, C., Fischl, B., Dale, A. & Halgren, E. Automatic parcellation of human cortical gyri and sulci using standard anatomical nomenclature. NeuroImage 53, 1–15 (2010).

91. Thomas Yeo, B. et al. The organization of the human cerebral cortex estimated by intrinsic functional connectivity. Journal of Neurophysiology 106, 1125–1165 (2011).

92. Fischl, B. et al. Automatically parcellating the human cerebral cortex. Cerebral Cortex 14, 11–22 (2004).

93. Hamilton, L. S., Chang, D. L., Lee, M. B. & Chang, E. F. Semi-automated anatomical labeling and inter-subject warping of high-density intracranial recording electrodes in electrocorticography. Frontiers in Neuroinformatics 11, 62 (2017).

94. Bastos, A. M. & Schoffelen, J.-M. A tutorial review of functional connectivity analysis methods and their interpretational pitfalls. Frontiers in Systems Neuroscience 9, 175 (2016).

95. Cao, J. & Worsley, K. The geometry of correlation fields with an application to functional connectivity of the brain. The Annals of Applied Probability 9, 1021–1057 (1999).

96. Fisher, R. A. Frequency distribution of the values of the correlation coefficient in samples from an indefinitely large population. Biometrika 10, 507–521 (1915).

97. Silver, N. C. & Dunlap, W. P. Averaging correlation coefficients: should Fisher’s z transformation be used? Journal of Applied Psychology 72, 146 (1987).

